# Biochemical and genetic dissection of the RNA-binding surface of the FinO domain of *Escherichia coli* ProQ

**DOI:** 10.1101/2023.04.25.538249

**Authors:** Ewa M. Stein, Suxuan Wang, Katherine Dailey, Chandra M Gravel, Shiying Wang, Mikołaj Olejniczak, Katherine E Berry

**Affiliations:** Institute of Molecular Biology and Biotechnology, Faculty of Biology, Adam Mickiewicz University, Uniwersytetu Poznańskiego 6, 61-614 Poznań, Poland; Program in Biochemistry, and Mount Holyoke College, South Hadley, MA, 01075, USA; Department of Chemistry, Mount Holyoke College, South Hadley, MA, 01075, USA

## Abstract

RNA-binding proteins play important roles in bacterial gene regulation through interactions with both coding and non-coding RNAs. ProQ is a FinO-domain protein that binds a large set of RNAs in *Escherichia coli*, though the details of how ProQ binds these RNAs remain unclear. In this study, we used a combination of *in vivo* and *in vitro* binding assays to confirm key structural features of *E. coli* ProQ’s FinO domain and explore its mechanism of RNA interactions. Using a bacterial three-hybrid assay, we performed forward genetic screens to confirm the importance of the concave face of ProQ in RNA binding. Using gel shift assays, we directly probed the contributions of ten amino acids on ProQ binding to seven RNA targets. Certain residues (R58, Y70, and R80) were found to be essential for binding of all seven RNAs, while substitutions of other residues (K54 and R62) caused more moderate binding defects. Interestingly, substitutions of two amino acids (K35, R69), which are evolutionarily variable but adjacent to conserved residues, showed varied effects on the binding of different RNAs; these may arise from the differing sequence context around each RNA’s terminator hairpin. Together, this work confirms many of the essential RNA-binding residues in ProQ initially identified *in vivo* and supports a model in which residues on the conserved concave face of the FinO domain such as R58, Y70 and R80 form the main RNA-binding site of *E. coli* ProQ, while additional contacts contribute to the binding of certain RNAs.

## INTRODUCTION

Small RNAs (sRNAs) participate in the regulation of gene expression in bacteria, contributing to the maintenance of cellular homeostasis, and adaptation to environmental changes (Wagner and Romby 2015; Gorski et al. 2017; Adams and Storz 2020). Hfq and ProQ are both global RNA-binding proteins involved in sRNA-dependent regulation in *Escherichia coli* and *Salmonella enterica*, which bind distinct, but partly overlapping RNA pools (Holmqvist et al. 2016; Melamed et al. 2016; Smirnov et al. 2016; Holmqvist et al. 2018; Melamed et al. 2020). ProQ binds hundreds of RNAs in *E. coli* and *S. enterica* (Holmqvist et al. 2018; Melamed et al. 2020), and contributes to such processes as DNA maintenance (Smirnov et al. 2016; Smirnov et al. 2017), adaptation to osmotic stress (Kerr et al. 2014; Melamed et al. 2020), motility (Rizvanovic et al. 2021), carbon source utilization (El Mouali et al. 2021b), adaptation to nutrient availability (Avrani et al. 2017; Knöppel et al. 2018; Gross et al. 2020; Katz et al. 2021), and virulence (Westermann et al. 2019). While Hfq has many well-established mechanisms of action (*e.g.* influencing sRNA-mRNA base pairing, sRNA lifetimes, mRNA translation, and RNA folding (Soper and Woodson 2008; Updegrove et al. 2016; Chen and Gottesman 2017; Andrade et al. 2018; Kavita et al. 2018)), much less is known about the detailed mechanisms used by ProQ to bind RNA and regulate gene expression.

ProQ belongs to the FinO family of proteins. Each member contains a conserved RNA-binding FinO domain, named after the founding member of this family, the F-like plasmid (Fʹ) FinO protein (Ghetu et al. 2000; Glover et al. 2015; Olejniczak and Storz 2017; Holmqvist et al. 2020) (Suppl. Figs S1 and S2). Proteins from this family are present in numerous proteobacteria (Attaiech et al. 2017). In *E. coli* and *S. enterica* ProQ, the N-terminal FinO domain (NTD) is connected via a long, positively charged linker to the C-terminal Tudor domain (CTD) (Gonzalez et al. 2017). This FinO domain has been shown to be the primary RNA binding site of ProQ *in vitro* and *in vivo* (Chaulk et al. 2011; Pandey et al. 2020; Stein et al. 2020).

Multiple approaches have been used to identify the residues of FinO-domain proteins that participate in RNA binding. For instance, the surfaces of the *E. coli* ProQ FinO domain that are protected by the binding of two RNAs, *cspE* and SraB, have been mapped using hydrogen-deuterium exchange experiments (Gonzalez et al. 2017). Crosslinking experiments in the homologous Fʹ-like FinO protein showed contacts between a FinP RNA fragment and several amino acid residues located mainly on the concave face of the FinO domain (Ghetu et al. 2002). The role of the concave face in RNA binding has also been supported by NMR studies of *Lp* Lpp1663 protein, which showed strong chemical shifts on the concave side of the FinO domain upon the binding of a U_6_ oligoribonucleotide or a hairpin derived from RaiZ sRNA (Immer et al. 2020). In addition, a recent co-crystal structure was solved showing how the terminator hairpin of *L. pneumophila* RocR binds to the FinO domain of RocC (Kim et al. 2022) (see discussion). This structure joins an additional four experimentally determined structures of FinO-domain orthologs from various bacterial species (Suppl. Fig S2; Ghetu et al. 2000; Chaulk et al. 2010; Gonzalez et al. 2017; Immer et al. 2020) and, more recently, AlphaFold predictions of these structures (Jumper et al. 2021).

We have previously utilized a bacterial-three hybrid (B3H) assay (Berry and Hochschild 2018; Stockert et al. 2022) to detect ProQ-RNA interactions (Pandey et al. 2020; Stein et al. 2020) and screen the effects of mutations in ProQ on RNA binding *in vivo*. This study was the first to examine the effects of amino-acid substitutions on the RNA-binding activity of *E. coli* ProQ. Both site-directed mutagenesis and unbiased screens converged on similar take-aways from this study, identifying a set of residues that contribute to SibB and *cspE* RNA binding by the FinO domain of ProQ (Pandey et al. 2020). The NMR-derived structural model available for this domain of *E. coli* ProQ (Gonzalez et al. 2017) suggested most of the RNA-binding residues fell on the concave surface, which had been implicated in RNA binding in studies performed on other FinO-domain homologs (Ghetu et al. 2000). Intriguingly, a single residue required for RNA interaction – the highly conserved arginine 80 – was modeled by the NMR structure (Gonzalez et al. 2017) to fall on the convex face of the NTD, opposite from the other RNA-binding residues on *E coli* ProQ and from the position of homologous residues in the solved structures of other FinO domain proteins (FinO, NMB1681, Lpp1663; (Ghetu et al. 2000; Chaulk et al. 2010; Gonzalez et al. 2017; Immer et al. 2020)) as well as its position on the AlphaFold structural prediction for *E. coli* ProQ (Jumper et al. 2021). The latter, however, is informed by homology so could be biased by the homologous protein structures if *E. coli* ProQ differed in this respect from other FinO domains. The location of this conserved and critical RNA-binding residue on the convex face raised the question of whether it constitutes an additional contact point in *E. coli* ProQ in addition to the concave surface.

At the end of our previous work, three primary questions remained unanswered: 1) how do we explain that the amino acid residues found to be required for RNA binding on the FinO domain of *E. coli* ProQ were quite distant from one another; 2) will the effects of mutations observed for the few RNAs examined so far hold true for a more diverse set of RNAs bound by ProQ; and 3) whether all of the critical residues for *in vivo* interactions are directly involved in RNA binding, rather than mediating indirect cellular effects?

In this work, we have used both *in vivo* and *in vitro* binding experiments to resolve these unanswered questions about ProQ-RNA interactions. We have probed the *in vivo* structure of the FinO domain of *E. coli* ProQ, using our B3H assay and a compensatory mutagenesis screen to shed light on the location of the conserved R80 residue. The results of this unbiased screen support a structural model in which the side chain of R80 points through to the concave face of the FinO domain, close to other RNA-binding residues and to the position of this side chain in structural homologs. Building on this structural insight, we have conducted the first *in vitro* site-directed mutagenesis experiments with *E. coli* ProQ to probe the contributions of both conserved and more variable amino acids on RNA binding. Gel-shift and quantitative analysis of twelve ProQ mutants revealed that certain amino acids have differential contributions to binding across seven natural RNA ligands, suggesting that ProQ may utilize distinct contacts with different RNA ligands. Further, we explored the properties of RNA structure that contributed to differential interactions with ProQ’s FinO domain and demonstrated the importance of single-stranded regions both upstream and downstream of terminator hairpins. Overall, this work advances a model in which the concave face of the FinO domain serves as the main RNA-binding site of *E. coli* ProQ and residues on the periphery of this surface tune interactions with different RNA ligands.

## RESULTS

### The concave-face pocket of FinO-domain is conserved in *E. coli* ProQ

We previously used an *in vivo* bacterial three-hybrid (B3H) assay to examine the effects of mutations on ProQ-RNA interactions (Fig. 1A; see methods for detailed description). One mutation that caused very strong defects in RNA interaction was R80A. To further explore the role arginine 80 plays in RNA binding, and whether its role goes beyond electrostatics, we created an R80K substitution, replacing arginine with the other basic residue lysine. We tested RNA interactions of both R80A and R80K ProQ variants using our B3H assay with three RNA ligands: *cspE*-3ʹ, SibB, and *malM*-3ʹ. Surprisingly, even the seemingly very conservative substitution of arginine to lysine at position 80 eliminated RNA binding to a similar extent as an alanine for all RNAs tested (Fig. 1B, C), even as western blotting showed that all three variants of α-ProQ were expressed to similar extents (Fig. 1C; Suppl. Fig S3).

**Figure 1.**
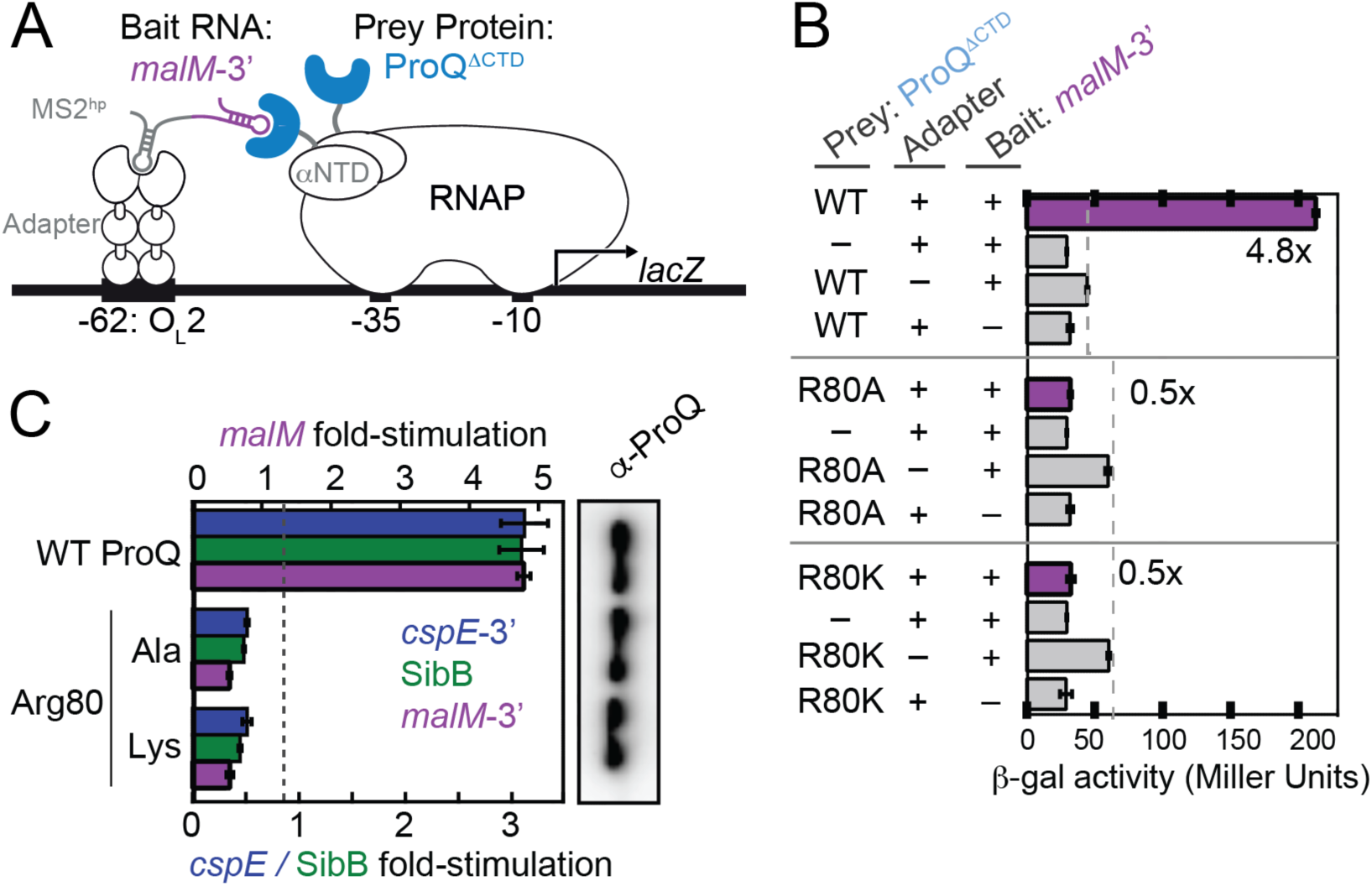
A positive charge at conserved position 80 is not sufficient for RNA interaction *in vivo* or *in vitro*. (A) Design of B3H system to detect interaction between ProQ and an RNA *(malM* 3ʹUTR). Interaction between the protein moiety and RNA moiety fused, respectively, to the NTD of the alpha subunit of RNAP (α) and to one copy of the MS2 RNA hairpin (MS2^hp^) activates transcription from test promoter, which directs transcription of a *lacZ* reporter gene (Berry and Hochschild 2018). Compatible plasmids direct the synthesis of the α-fusion protein (under the control of an IPTG-inducible promoter), the CI-MS2^CP^ adapter protein (under the control of a constitutive promoter; pCW17) and the hybrid RNA (under the control of an arabinose-inducible promoter). (B) Results of β-galactosidase assays performed in Δ*hfq* p*lac*-O_L_2–62 reporter strain cells containing three compatible plasmids: one (α-ProQ) that encoded α (–) or the α-ProQ^ΔCTD^ (pKB955; residues 2-176) fusion protein (WT or an R80A or R80K mutant), another (CI-MS2^CP^) that encoded λCI (–) or the λCI-MS2^CP^ fusion protein (+), and a third (Bait) *c*) that encoded a hybrid RNA with the 3ʹ UTR of *malM* (pKB1210) following one copy of an MS2^hp^ moiety (+) or an RNA that contained only the MS2^hp^ moiety (–). Cells were grown in the presence of 0.2% arabinose and 50 μM IPTG (see Methods). Bar graphs show the averages of three independent measurements and standard deviations. (C) (left) Results of B3H assays detecting interactions between α-ProQ^ΔCTD^ and three RNA baits. β-galactosidase assays were performed as in (B) but with three hybrid RNA constructs (MS2^hp^-*malM*-3ʹ, MS2^hp^-*cspE*-3ʹ, MS2^hp^-SibB). The bar graph shows the fold-stimulation over basal levels as averages and standard deviations of values collected from three independent experiments conducted in triplicate across multiple days. (right) Western blot to compare steady-state expression levels of mutant α-ProQ^ΔCTD^ fusion proteins. Lysates were taken from the corresponding β-gal experiment containing MS2^hp^-*malM-*3ʹ and all other hybrid components at 50 μM IPTG. Following electrophoresis and transfer, membranes were probed with anti-ProQ antibody.

The inability of a lysine to substitute for arginine at residue 80 is analogous to the strongly deleterious effect we previously observed of a conservative Y70F substitution on RNA binding *in vivo* (Pandey et al. 2020). We wondered whether the positive charge and aromatic ring were not the critical feature of this particular arginine and tyrosine, respectively. To explicitly test whether any other amino acid could support RNA binding at either position 70 or 80 in the FinO domain of *E. coli* ProQ, we constructed saturation mutagenesis libraries at each of these positions and screened for colonies showing any amount of blue over negative controls with *malM*-3ʹ RNA as bait in the B3H assay. Sequencing results of isolated plasmids showed that all codons encoding arginine at position 80 and tyrosine at position 70 were recovered (Suppl. Table S1), but no other substitutions at either position were found to support detectable binding *in vivo*. This confirms that both Y70 and R80 are uniquely required for RNA interaction. Indeed, these residues are both highly conserved across FinO-domain sequences and are located near one another in a concave pocket in most solved structures of FinO-domains (Suppl. Figs. S1 and S2).

Given that both Y70 and R80 were important for *E. coli* ProQ’s interaction with RNA, it was important to clarify if these residues are indeed located on opposite faces of the protein as previously suggested (Gonzalez et al. 2017; Pandey et al. 2020), especially because these residues are found on the same face of the FinO domain in the structures of other homologs (Suppl. Fig. S2) (Ghetu et al. 2000; Chaulk et al. 2010; Immer et al. 2020; Kim et al. 2022). We reasoned that the chemical change in ProQ^R80K^ was minor enough that it should be possible to rescue with compensatory mutations of nearby residues – and that the location of these compensatory substitutions would provide evidence regarding the likely structural position for R80 in the structure of *E. coli* ProQ found *in vivo*. To create compensatory mutations that could rescue RNA binding of ProQ^R80K^, we selected three regions of the primary sequence that were close to proposed positions of R80 in *E. coli* ProQ (convex face) or in other FinO domain proteins (concave face; Fig. 2A and Suppl. Fig. S2). We then used a saturation-mutagenesis strategy (Suppl. Fig. S4A) to create a library of all possible single point-mutations at each of these amino-acid positions on the pPrey-ProQ^R80K^ plasmid.

**Figure 2.**
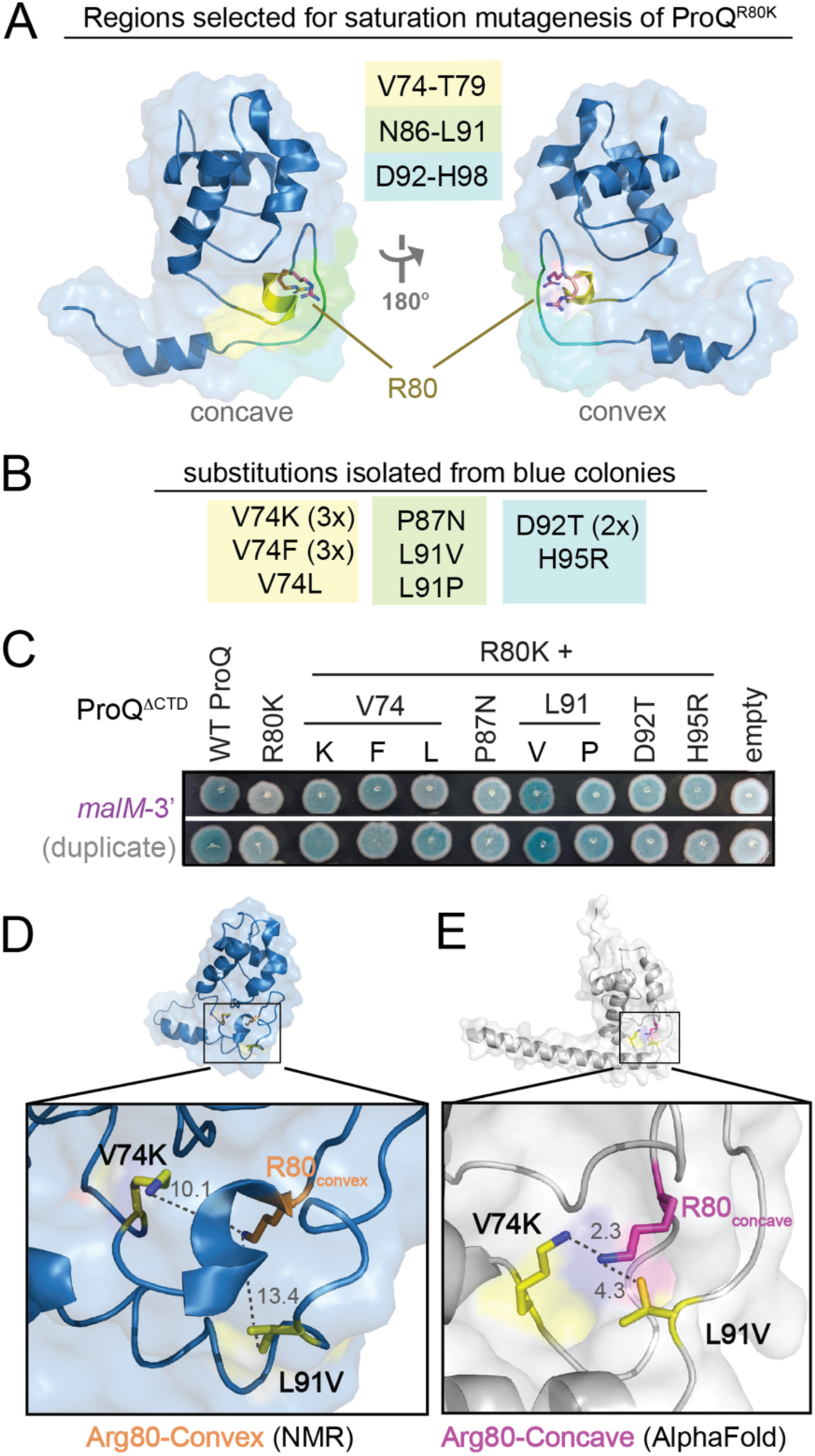
Saturation mutagenesis screen to identify compensatory mutants that rescue RNA-binding activity of ProQ^R80K^. (A) Residues in three regions were selected for mutagenesis. These regions are highlighted in yellow, green and cyan on the NMR structure of the ProQ NTD (Gonzalez et al. 2017). Arginine 80 is shown in orange sticks in the convex-face position found in the NMR structure and in pink sticks in a hypothetical concave-face position that would be analogous to its position in other orthologs. Saturation mutagenesis libraries were created from pPrey-ProQ^R80K^, in which the sequence one codon at a time was varied (see Methods and Suppl. Fig. S4A) (B) Results from compensatory mutant screen. pPrey-ProQ^R80K^ plasmids that produced blue colonies on X-gal-indicator plates in OL2-62-l*acZ* reporter cells in the presence of pBait-*malM-*3ʹ and pAdapter were sequenced, and the substitutions encoded by mutated codons are listed. A subset of these codons were identified in multiple sequenced colonies, indicated by 2x and 3x. (C) Results of B3H assays confirming the effects of compensatory mutants in a plate-based B3H assay, detecting interactions between variants of ProQ^ΔCTD^ with *malM-*3ʹ RNA. β-gal assays were performed with *Δhfq* reporter-strain cells containing three compatible plasmids: one that encoded the CI-MS2^CP^ fusion protein, another that encoded α or an α-ProQ^ΔCTD^ fusion protein (wild type, WT, or the indicated mutant), and a third that encoded a hybrid RNA (MS2^hp^-*malM*-3ʹ). Cells were grown in the presence of 0.2% arabinose and 50 μM IPTG and then plated on X-gal indicator plates with selective antibiotics (see methods). (D,E) Positions of two strongest compensatory substitutions (V74 and L91) are shown on (D) the NMR structure (PDB ID 5NB9, (Gonzalez et al. 2017)) and (E) the Alphafold Model (Varadi et al. 2022) of the FinO domain of *E. coli* ProQ. Distances between the terminal atom of each side chain (R80K, V74K, L71V) are shown in Angstroms and visualized with dashed lined. The side chains were mutated in PyMol to show predicted structures of the compensatory substitutions.

To screen for compensatory mutations that could partially rescue RNA binding of ProQ^R80K^, saturation mutagenesis libraries were transformed into B3H reporter cells containing pAdapter and pBait-*malM* plasmids. Transformants were plated on X-gal indicator plates, and plasmids were purified and sequenced from colonies that were more blue than a pPrey-empty control. Mutations that partially rescued ProQ-*malM* binding were identified from each of the three mutagenesis libraries constructed (Fig. 2B); multiple mutations were identified at the same amino acid positions (V74 and L91) and several of the mutant plasmids were isolated and sequenced multiple times. When isolated plasmids were re-transformed into fresh B3H reporter cells, six of the eight initial hits showed above-background interaction with *malM*-3ʹ RNA in qualitative plate-based assays (Fig. 2C). The strongest of these substitutions (L91V and V74K) also resulted in modest rescue of ProQ^R80K^ binding to *malM-3*ʹ in liquid β-gal assays and to *cspE*-3ʹ and SibB RNAs in plate-based assays (Suppl. Fig. S4B, C). The fact that compensatory effects of substitutions on RNA binding were more evident in plate-based assays with *malM*-3ʹ than in liquid β-gal assays or interactions with other RNA ligands is not surprising since the screen in which they were identified was conducted in a plate-based screen with *malM*-3ʹ RNA.

We next mapped the positions of the strongest confirmed compensatory substitutions to two available structures for the ProQ^NTD^ (Gonzalez et al. 2017; Jumper et al. 2021; Varadi et al. 2022). While the two models share an overall fold of the FinO domain, they differ in the region containing R80. While V74 and L91, the sites of the strongest compensatory mutations, are 10-13Å away from the terminal atom of a convex-facing R80 in the NMR structure (Fig. 2D) (Gonzalez et al. 2017), these residues are much closer (2-4Å) in the AlphaFold structural model, which places R80 on the concave face (Fig. 2E) (Jumper et al. 2021; Varadi et al. 2022). Indeed, the AlphaFold model rationalizes why the conservative R80K would be such a deleterious substitution: lysine’s shorter aliphatic side would pull the terminal polar/charged functional group into the hydrophobic core of the protein. The model also explains why an V74K substitution would rescue a lysine at position 80, as it provides a second amine in hydrogen-bonding distance of R80K in an otherwise hydrophobic environment (Fig. 2E).

Together, these results lend experimental validation to the AlphaFold model of *E. coli* ProQ’s FinO domain, in which all critical RNA-binding residues, including R80, are located on the concave face of the FinO domain (Fig. 2E, Suppl. Fig S2). In this model, the concave face of the FinO domain looks quite like other structural homologs, with the highly conserved Y70 and R80 residues positioned near one another in a concave pocket.

### Central concave-pocket residues are essential for RNA binding *in vitro*

With this refined structural model for *E. coli* ProQ^NTD^ in mind, we wished to further explore how specific FinO-domain residues contribute to RNA binding. In particular, we wanted to test whether conclusions from B3H studies with *cspE*-3ʹ and SibB RNAs about the importance of specific residues on RNA binding (Pandey et al. 2020) would also hold true with purified components and for other RNA ligands of ProQ. For this purpose, we compared RNA binding to ProQ^NTD^ mutants using electrophoretic mobility shift assays, as previously conducted (Stein et al. 2020). We selected seven RNAs for this study, each of which had been found to bind to ProQ *in vivo* (Suppl. Fig S5; (Holmqvist et al. 2018; Melamed et al. 2020)). These RNAs included *malM*-3ʹ, the top RNA ligand identified by RIL-seq of ProQ (Melamed et al. 2020), two versions of *cspE* 3ʹ-UTR (a 52-nt fragment (*cspE*-3ʹ) and an 81-nt fragment (*cspE*81-3ʹ)), *gapA*-3ʹ and SibA, all of which are specific ligands of ProQ *in vivo*, and RybB, which is bound by both ProQ and Hfq *in vivo*. With the exceptions of SibB and RybB, the binding of all these RNAs had already been studied with purified ProQ^NTD^ (Stein et al. 2020).

For binding studies, we used a 130-aa long version of ProQ’s NTD, the RNA binding properties of which had been previously studied (Chaulk et al. 2011; Pandey et al. 2020; Stein et al. 2020). We first compared the binding affinities of the seven RNA ligands with WT ProQ^NTD^. The observed *K*_d_ values for ProQ^NTD^ binding to *cspE*-3ʹ, *cspE*81-3ʹ, *malM*-3ʹ, SibA, and SibB fell within in a similar range (4 to 10 nM); only *gapA*-3ʹ, and RybB bound weaker with *K*_d_ values of 14 nM and 26 nM, respectively (Figs. 3, 4, Table 1, Suppl. Figs. S6-S12). Overall, the seven RNAs bound WT ProQ^NTD^ with a relatively narrow range of affinities.

**Figure 3.**
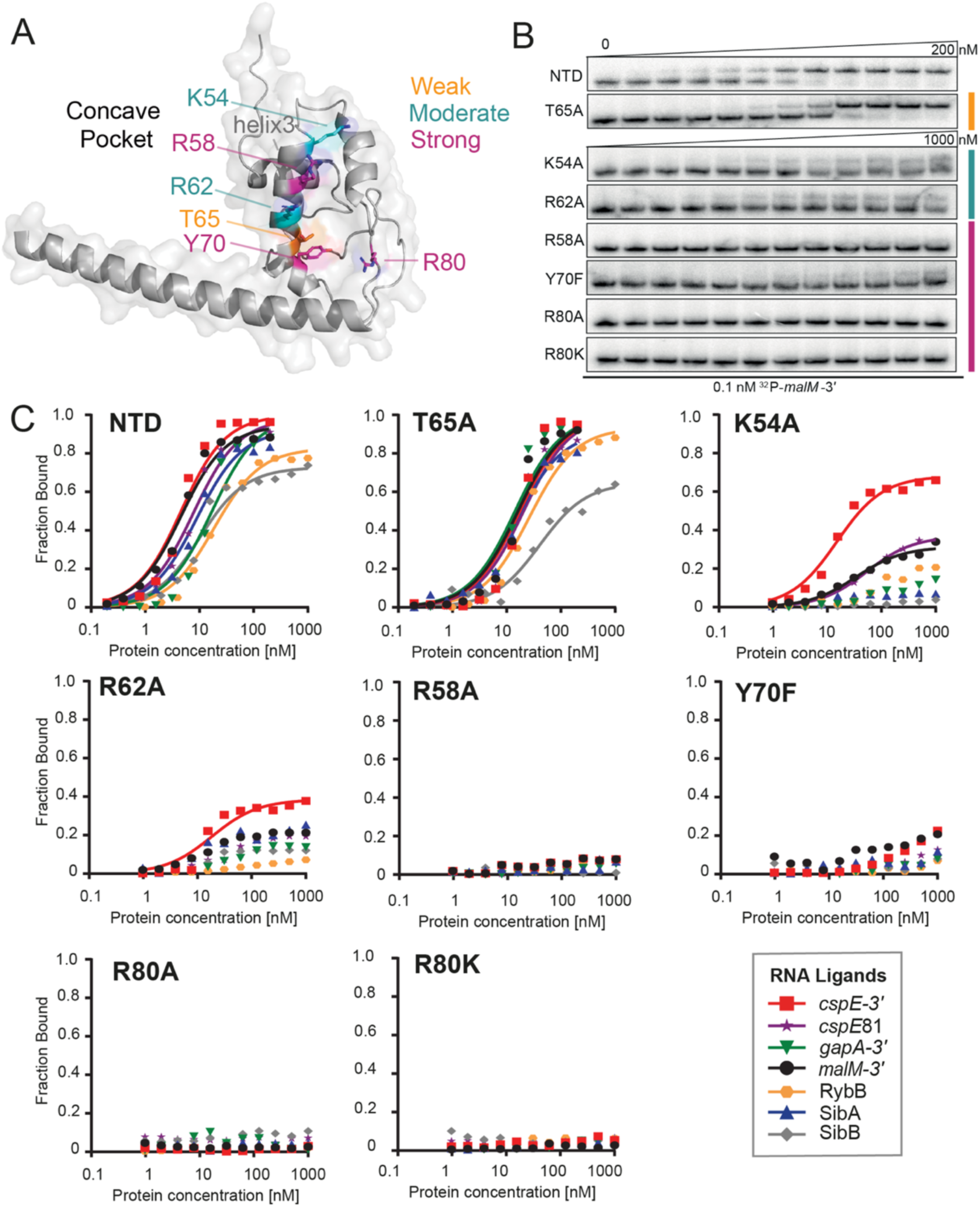
Mutations in the central pocket have strong detrimental effects on the binding of seven RNAs. (A) The location of substitutions on the structure of ProQ^NTD^ (Varadi et al. 2022). (B) The raw gel data for 1 nM ^32^P-labeled *malM*-3’ binding to the 8 proteins. (C) The plots of fraction bound versus protein concentration for the binding of *cspE*-3’, *cspE*81-3ʹ, *gapA*-3’, *malM*-3’, RybB, SibA and SibB RNAs to WT ProQ^NTD^, and T65A, K54A, R62A, R58A, Y70F, R80A, and R80K mutants measured using the gelshift assay. The data sets in which maximum fraction bound was above 40% were analyzed by fitting to the quadratic equation. Raw gel data for all RNAs are presented in Suppl. Figures S6-S12. The obtained *K*_d_ values are shown in Table 1.

**Table 1.**
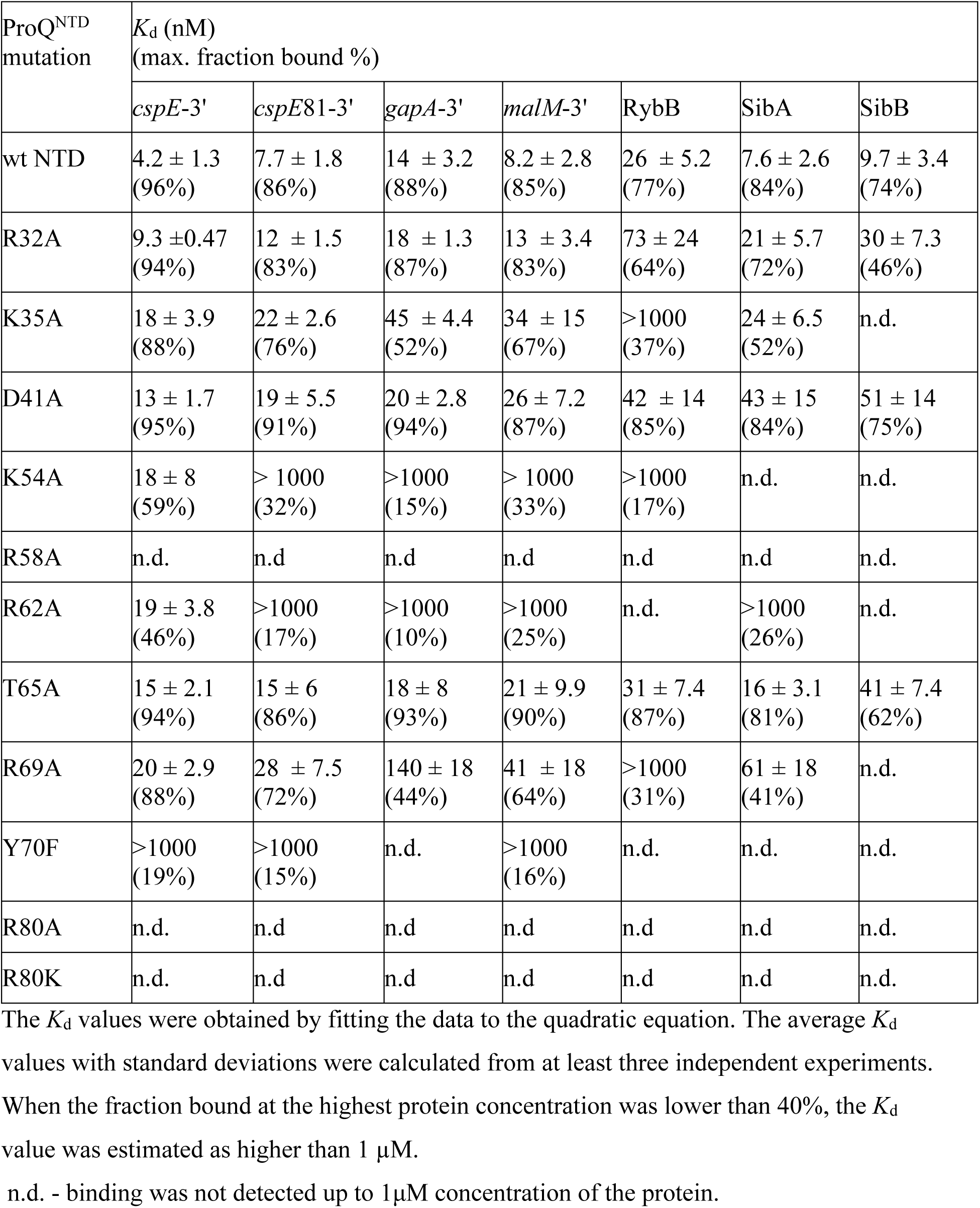
Equilibrium RNA binding to WT FinO domain and its mutants.

| ProQ<sup>NTD</sup><br>mutation | $K_d$ (nM)<br>(max. fraction bound %) | | | | | | |
| --- | --- | --- | --- | --- | --- | --- | --- |
|  | *cspE*-3' | *cspE*81-3' | *gapA*-3' | *malM*-3' | RybB | SibA | SibB |
| wt NTD | 4.2 ± 1.3<br>(96%) | 7.7 ± 1.8<br>(86%) | 14 ± 3.2<br>(88%) | 8.2 ± 2.8<br>(85%) | 26 ± 5.2<br>(77%) | 7.6 ± 2.6<br>(84%) | 9.7 ± 3.4<br>(74%) |
| R32A | 9.3 ± 0.47<br>(94%) | 12 ± 1.5<br>(83%) | 18 ± 1.3<br>(87%) | 13 ± 3.4<br>(83%) | 73 ± 24<br>(64%) | 21 ± 5.7<br>(72%) | 30 ± 7.3<br>(46%) |
| K35A | 18 ± 3.9<br>(88%) | 22 ± 2.6<br>(76%) | 45 ± 4.4<br>(52%) | 34 ± 15<br>(67%) | >1000<br>(37%) | 24 ± 6.5<br>(52%) | n.d. |
| D41A | 13 ± 1.7<br>(95%) | 19 ± 5.5<br>(91%) | 20 ± 2.8<br>(94%) | 26 ± 7.2<br>(87%) | 42 ± 14<br>(85%) | 43 ± 15<br>(84%) | 51 ± 14<br>(75%) |
| K54A | 18 ± 8<br>(59%) | > 1000<br>(32%) | >1000<br>(15%) | > 1000<br>(33%) | >1000<br>(17%) | n.d. | n.d. |
| R58A | n.d. | n.d. | n.d. | n.d. | n.d. | n.d. | n.d. |
| R62A | 19 ± 3.8<br>(46%) | >1000<br>(17%) | >1000<br>(10%) | >1000<br>(25%) | n.d. | >1000<br>(26%) | n.d. |
| T65A | 15 ± 2.1<br>(94%) | 15 ± 6<br>(86%) | 18 ± 8<br>(93%) | 21 ± 9.9<br>(90%) | 31 ± 7.4<br>(87%) | 16 ± 3.1<br>(81%) | 41 ± 7.4<br>(62%) |
| R69A | 20 ± 2.9<br>(88%) | 28 ± 7.5<br>(72%) | 140 ± 18<br>(44%) | 41 ± 18<br>(64%) | >1000<br>(31%) | 61 ± 18<br>(41%) | n.d. |
| Y70F | >1000<br>(19%) | >1000<br>(15%) | n.d. | >1000<br>(16%) | n.d. | n.d. | n.d. |
| R80A | n.d. | n.d. | n.d. | n.d. | n.d. | n.d. | n.d. |
| R80K | n.d. | n.d. | n.d. | n.d. | n.d. | n.d. | n.d. |

The $K_d$ values were obtained by fitting the data to the quadratic equation. The average $K_d$ values with standard deviations were calculated from at least three independent experiments. When the fraction bound at the highest protein concentration was lower than 40%, the $K_d$ value was estimated as higher than 1 μM.
n.d. - binding was not detected up to 1 μM concentration of the protein.

Because the *in vivo* B3H data showed the importance of universally conserved residues tyrosine 70 and arginine 80 for the binding of *cspE-*3ʹ, *malM* 3ʹ and SibB (Figs. 1, 2; (Pandey et al. 2020), we next compared the effects of mutations at these positions on the binding of the seven RNAs. The Y70F mutant had a dramatic effect on RNA binding across all RNAs, with the fraction of RNA bound reaching no higher than 20% at the highest concentration of ProQ^NTD^ (1 μM) for *cspE*-3ʹ, *cspE*81-3ʹ, and *malM*-3ʹ RNAs, and no binding detected at all for *gapA*-3ʹ, RybB, SibA, and SibB RNAs (Fig. 3, Table 1, Suppl. Figs. S6-S12). Because this and subsequent mutations affected the maximum fraction of RNA bound to ProQ, Table 1 reports the maximum fraction bound for each mutant as well as *K*_d_ values for experiments where the fraction bound reached at least 40%. Substitutions at R80 were even more deleterious than those at Y70: no binding was detected for any of the seven RNAs to either R80A or R80K mutants (Fig. 3B,C, Table 1, Suppl. Figs S6-S12), consistent with the strong effect of both mutants *in vivo* (Fig. 1C). The fact that substitutions of Y70 and R80 were so deleterious for the binding of all seven RNAs supports the conclusion that these amino acid residues are universally important for the binding of natural RNA ligands of ProQ.

Next, we analyzed the role of threonine 65, which is close to Y70 on the concave face of the FinO domain (Fig. 3A). While T65 was not identified in our *in vivo* mutagenesis screens (Pandey et al. 2020), we hypothesized that it could be important for RNA binding based on its proximity to the side chain of Y70 as well as its conservation as a threonine or a structurally similar serine or cysteine (Suppl. Fig. S1). Despite these factors, the T65A mutation had rather small effects on RNA binding by ProQ^NTD^, ranging from no effect on the binding of *gapA*-3ʹ and RybB, to 2-fold weaker binding than WT ProQ^NTD^ for *cspE*81-3ʹ, *malM*-3ʹ, and SibA, to 4-fold weaker binding for *cspE*-3ʹ and SibB (Fig. 3B, C, Table 1, Suppl. Figs S6-S12). In keeping with the modest effects on *K*_d_, the T65A mutant did not alter the maximum fraction bound of any RNA compared to WT ProQ^NTD^. These data suggest that T65 makes only small contributions to RNA binding despite its partial conservation and positioning adjacent to the essential Y70 residue.

Arginine 62 is located an additional helical turn away from T65 in helix H3 (Fig. 3A). We were intrigued that the identity of R62 is not evolutionarily conserved, despite its positive charge and location in the concave-face pocket of ProQ’s FinO domain. R62 therefore seemed a good candidate for an RNA contact that could be unique to *E. coli* ProQ relative to other FinO-domain proteins. Indeed, gel-shift data showed that the R62A mutation had a strongly detrimental effect on the binding of all seven RNAs, as the maximum fraction bound was below 40% for most RNAs (Fig. 3, Table 1, Suppl. Figs. S6-S12). The mutant protein’s interaction with *cspE*-3ʹ was the strongest of all the RNAs with a *K*_d_ value only 4-fold weaker than to WT ProQ^NTD^, although with a fraction bound just above 40% at the highest concentration tested of R62A ProQ^NTD^ (Table 1). The maximum fractions bound observed for other RNAs were lower still – around 25% for *malM*-3ʹ and SibA, and below 20% for *cspE*81-3ʹ and *gapA*-3ʹ – while no binding was detected for RybB and SibB. Although there seems to be a range of effects of R62A on different RNAs, these differential effects are impossible to precisely quantify because of the overall strongly weakened binding. Overall, the R62A mutation is strongly detrimental to binding of all seven RNAs, although not to the same degree as mutations of Y70 or R80.

We next turned our attention to two basic residues that were found to be important for *in vivo* ProQ-RNA interactions and were hypothesized to interact with the double-helical portion of a terminator hairpin (Pandey et al. 2020). These residues – arginine 58 and lysine 54 – are located one or two additional helical turn(s) further along H3 from R62 on the concave face of ProQ (Fig. 3A). We first analyzed the effects of an alanine substitution of the R58 residue, the closer of the two residues to Y70, and found that an R58A mutation was strongly deleterious for binding of all seven RNAs. As with R80A, no binding of any RNA was detected up to concentrations of 1 μM of the R58A mutant (Fig. 3, Table 1, Suppl. Figs. S6-S12), demonstrating an essential role of R58 in RNA binding by the ProQ FinO domain. On the other hand, the effects of a K54A mutation were more moderate: the maximum fraction bound only reached about 60% for *cspE*-3ʹ, 30% for *malM*-3ʹ and *cspE*81-3ʹ, and < 20% for *gapA*-3ʹ and RybB, while no binding was detected for SibA and SibB (Fig. 3, Table 1, Suppl. Figs S6-S12). These effects are comparable to those of the R62A mutation described above. Together, these data show that residues K54 and R58 on helix H3 of the FinO domain play important roles in RNA binding.

### RNA ligands differ in their sensitivity to amino-acid substitutions on the periphery

In addition to the central part of the concave-face pocket explored above, our previous B3H studies indicated that some residues on the periphery of the concave face, such as lysine 35 and aspartate 41, could also contribute to RNA binding (Pandey et al. 2020). Interestingly, B3H studies showed that the K35A mutation had more detrimental influence on the binding of SibB than *cspE*, which suggests that K35 could contribute differentially to binding of distinct RNAs (Pandey et al. 2020). When the binding of our panel of seven RNAs was tested *in vitro* with the K35A ProQ^NTD^ mutant, there were indeed large differences in the maximum fraction bound reached by each RNA. While the maximum fraction of *cspE*-3ʹ and *cspE*81-3ʹ bound to the K35A mutant remained similar to that of WT ProQ, the K35A mutant only reached about 70% RNA bound with *malM*-3ʹ, and 35-50% with *gapA*-3ʹ, SibA and RybB. The effects on SibB were the strongest, with no binding detected up to 1 μM protein. For several RNAs that reached a much lower fraction bound with K35A than for WT ProQ, the *K*_d_ values calculated from the fits were still within 3-4-fold of the *K*_d_ for WT ProQ. That the differences in binding were more apparent from the maximum fractions bound rather than the *K*_d_ values could suggest that complexes formed by the K35A mutant with some RNAs were less stable during electrophoresis in native gels, leading to the underestimation of calculated *K*_d_ values. Alternatively, such a result could be explained if tight binding of some RNAs required a conformational change that was only efficiently induced by ProQ^NTD^ in the presence of K35. Importantly, the relative effects of the K35A mutation on the binding of *cspE*-3ʹ and SibB are in agreement with the results obtained by the B3H assay *in vivo*, which also showed a stronger negative effect of this mutation on the binding of SibB (Pandey et al. 2020).

We next examined effects of mutating aspartate 41, another residue located at the base of helix H2 (Fig. 4A) that was suggested by previous B3H results to be important for RNA binding (Pandey et al. 2020). *In vitro* binding data showed that the D41A substitution does not affect the *K*_d_ value of ProQ^NTD^ for *gapA*-3ʹ, but does have a detrimental effect on the binding affinity for other RNAs; the size of its effect ranged from ∼2-fold (*cspE*81-3ʹ and RybB) or 3-fold (*cspE*-3ʹ and *malM*-3ʹ) to more than 5-fold in the strongest cases (SibA and SibB) (Fig. 4, Table 1, Suppl. Figs. S6-S12). Interestingly, the binding of all seven RNAs to the D41A mutant reached the same maximum fraction bound as with WT ProQ^NTD^ (Table 1), in contrast to the effects of K35A above. In summary, both D41 and K35 contribute modestly to RNA binding by ProQ *in vitro*, with K35 having more varied effects across RNAs, perhaps mediated through kinetics or facilitating conformational changes in the RNA.

**Fig 4.**
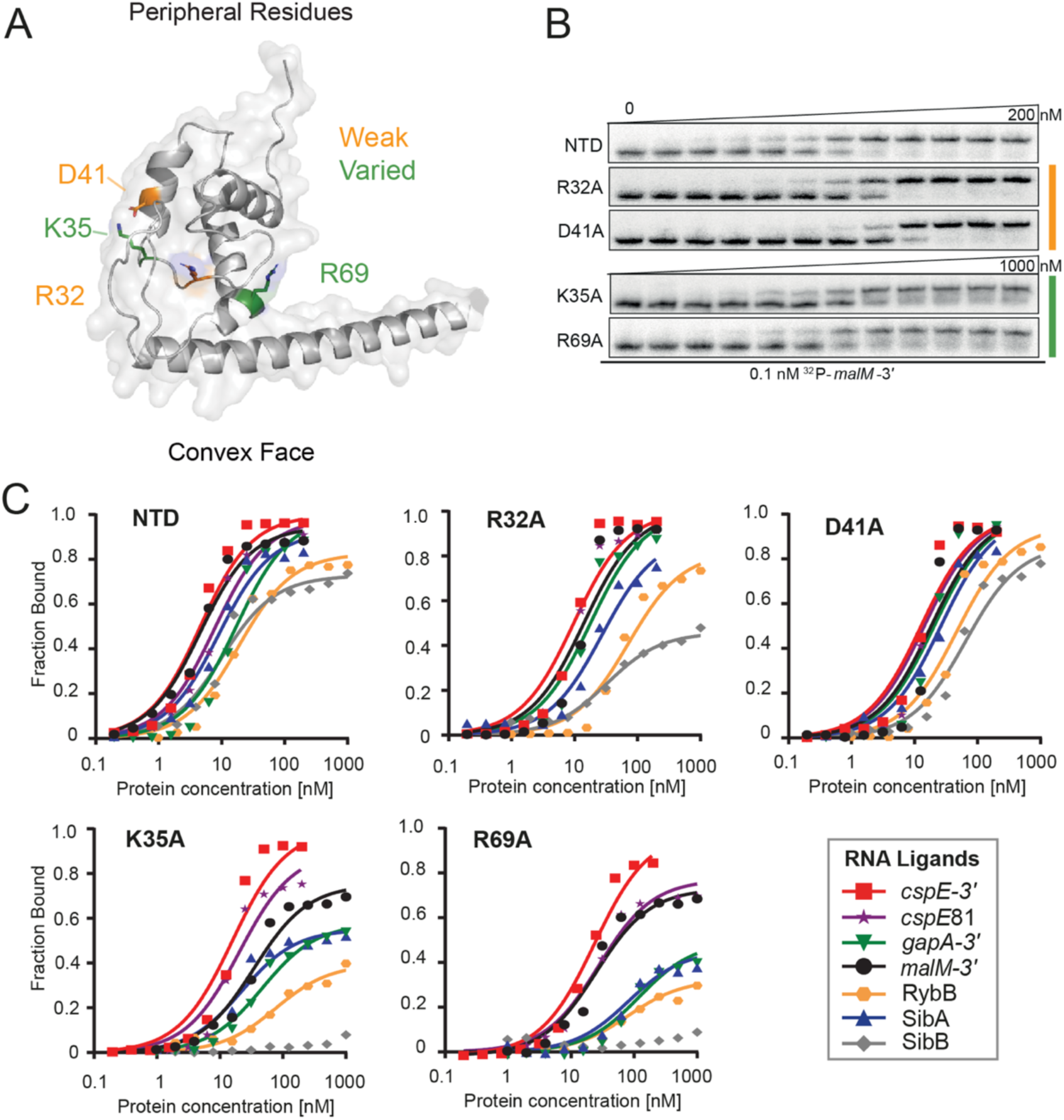
Mutations on the periphery of the central pocket have differential effects on the binding of seven RNAs. (A) The location of substitutions on the structure of ProQ^NTD^ (Varadi et al. 2022). (B) The raw gel data for 1 nM ^32^P-labeled *malM*-3’ binding to the 5 proteins. (C) The plots of fraction bound versus protein concentration for the binding of *cspE*-3’, *cspE*81-3ʹ, *gapA*-3’, *malM*-3’, RybB, SibA and SibB RNAs to WT ProQ^NTD^, and R32A, D41A, K35A, and R69A mutants measured using the gelshift assay. The data sets in which maximum fraction bound was above 40% were analyzed by fitting to the quadratic equation. Raw gel data for all RNAs are presented in Suppl. Figures S6-S12. The obtained *K*_d_ values are shown in Table 1.

Next, we moved further away from the concave-face pocket of ProQ, and asked whether positively charged amino acids on the convex face of the FinO domain could also affect RNA binding. We first examined the effects of arginine 32, which is conserved either as arginine or lysine in other FinO-domain proteins (Suppl. Fig. S1). *In vitro* binding data showed that an R32A mutation does not affect the *K*_d_ value of ProQ binding to *gapA*-3ʹ and *malM*-3ʹ and has only modest 2-to 3-fold detrimental effects on the binding of *cspE*-3ʹ, *cspE*81-3ʹ, RybB and SibA (Fig. 4, Table 1, Suppl. Figs. S6-S12). The only RNA that was markedly affected by this mutation was SibB; while its *K*_d_ value was only weakened ∼3-fold by the R32A mutation, its maximum fraction bound was reduced to just above 40% (Table 1). The fact that the R32A substitution has only small effects on the binding of most of the RNAs tested suggests that R32 does not make a universal key contact with RNA.

Another basic side chain exposed on the convex face of the FinO domain is arginine 69. Unlike R32, R69 is in an evolutionarily variable position, but its location next to Y70 in primary amino acid sequence suggested it could affect RNA binding (Suppl. Fig. S1). *In vitro* binding data showed that an R69A substitution had a 3-to 5-fold detrimental effect on the binding of *malM*-3ʹ and *cspE-*3ʹ, and an 8-to 10-fold effect on interactions with SibA and *gapA*-3ʹ (Fig. 4, Table 1, Suppl. Figs. S6-S12). The R69A mutation had an even stronger detrimental effect on the binding of other RNAs: the maximum fraction bound reached by RybB was below 40%, while the binding of SibB was not detected. The large range of effects of the R69A mutation were reminiscent of the effects of K35A, though perhaps qualitatively slightly stronger for each RNA. It is interesting to note that the rank-order of effects caused by K35A and R69A mutations on the maximum fraction bound reached by each RNA were similar, with *cspE*-3ʹ being the least affected by either mutation and SibB being the most affected (Table 1).

### The terminator hairpin drives the differential dependence of *cspE*-3′ and SibB on R69

The results above indicated that several residues, especially K35 and R69, made differential contributions to the binding of the seven RNAs we tested. To explore the basis for these differences, we focused on *cspE*-3ʹ and SibB RNAs, which fell on the opposite ends of the spectrum of susceptibility to ProQ^NTD^ mutants. Because the FinO domain of ProQ is known to specifically recognize the intrinsic terminators of its RNA ligands (Stein et al. 2020), we focused on this region of *cspE*-3ʹ and SibB. The two most striking differences between the terminator regions in these two RNAs are that SibB possesses a shorter oligoU tail that ends with 2 cytidines and that the base of *cspE*’s terminator hairpin contains several A-U base-pairs missing from SibB (Fig. 5). We introduced changes into SibB, the weaker binding RNA, to make its 3ʹ terminator more similar to *cspE*-3ʹ and compared the binding of these mutant RNAs to WT and R69A ProQ^NTD^. The first SibB mutant (SibB-5U) replaced the two terminal cytidines with uridines (Fig. 5A, B), resulting in an RNA with a total of five U’s in the tail, four of which are predicted to be single-stranded. While the SibB-5U mutation had no effect on the binding of WT ProQ^NTD^, it improved binding to the R69A mutant, with the mutant protein reaching a fraction bound of >30% when binding was undetectable for unmutated SibB RNA (Fig. 5B, Table 2, Suppl. Fig. S13). We next introduced additional mutations to make the double-stranded base of the SibB terminator hairpin more like that of *cspE*-3ʹ. This SibB-*cspE* chimera now had 4 A-U base pairs at the base of the terminator and an extended 8-uridine 3ʹ tail to match that of *cspE*-3ʹ (Fig. 5C, D). While the mutations in the SibB-*cspE* chimera had only slight effects on the binding of WT ProQ^NTD^, they were sufficient to substantially restore RNA binding to the R69A mutant, which reached a fraction bound of 60% with a *K*_d_ value of 31 nM (Fig. 5C, Table 2, Suppl. Fig. S13). These are considerable improvements from the original SibB RNA, which showed no detectable binding to the R69A mutant, though do not represent as strong of binding as seen with *cspE*-3ʹ RNA, which reached a maximum fraction bound of 88% to the R69A mutant with a *K*_d_ of 20 nM.

**Fig. 5.**
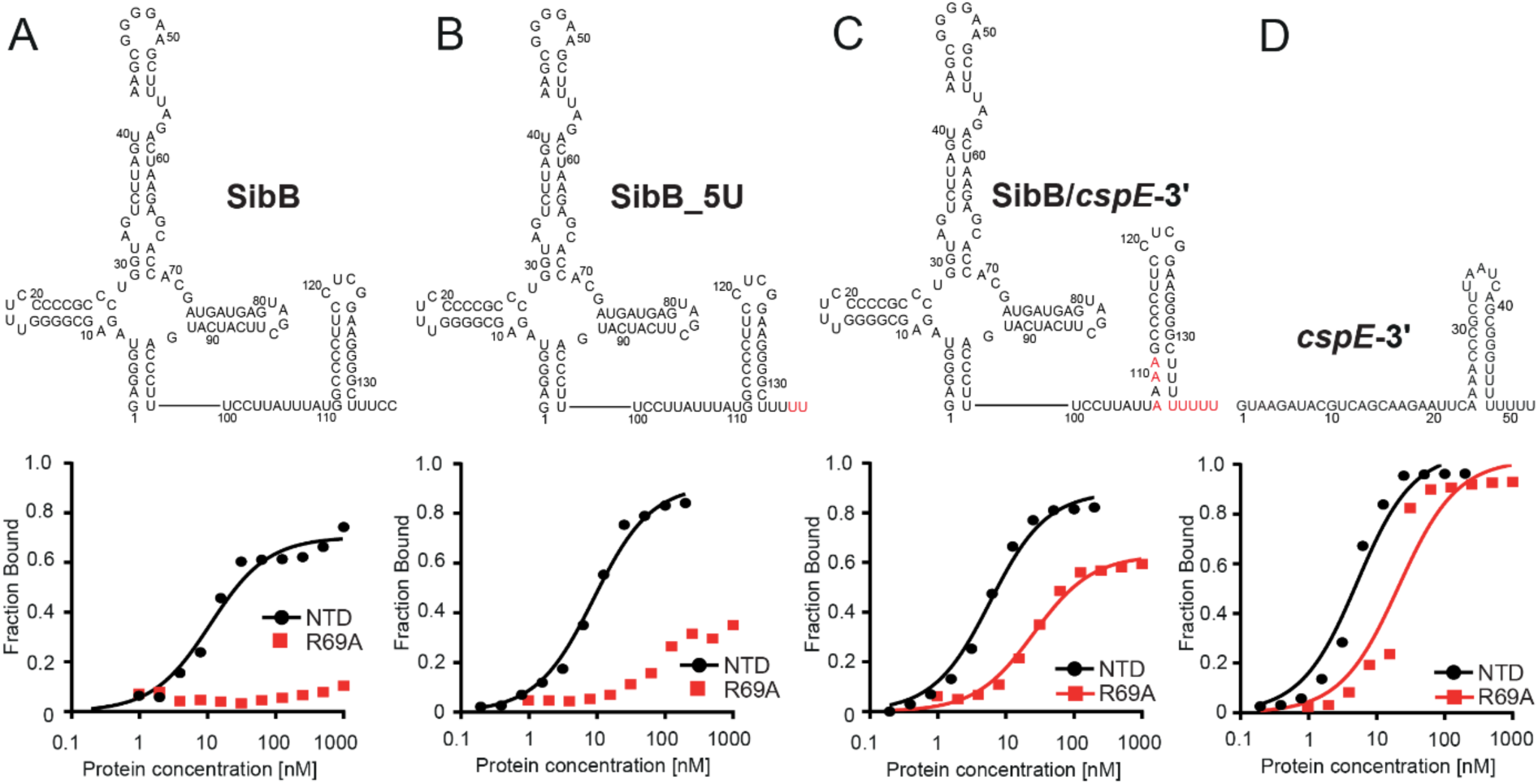
The sequence and structure of the lower part of the terminator hairpins explains differences in the binding of ProQ^NTD^ and its mutants by *cspE*-3’ and SibB RNAs. (A) – (D), The secondary structures of RNA molecules are shown above the corresponding plots of fraction bound data for WT ProQ^NTD^ (NTD) or its R69A mutant. The RNA secondary structures were predicted using *RNAStructure* software. The changes introduced into the sequence of SibB are marked in red font. The data sets in which maximum fraction bound was above 40% were analyzed by fitting to the quadratic equation. Data for SibB and *cspE*-3ʹ are the same as in Fig. 3. The obtained *K*_d_ values are shown in Table 2, and raw gel data are presented in the Suppl. Fig. S13.

**Table 2.** Mutations in the terminator hairpin of SibB improve its binding to the R69A mutant of ProQ^NTD^.

| ProQ<sup>NTD</sup><br>mutation | $K_d$ (nM)<br>(max. fraction bound, %) | | | |
| --- | --- | --- | --- | --- |
|  | SibB | SibB_5U | SibB/ *cspE*-3' | *cspE*-3' |
| wt NTD | 9.7 ± 3.4<br>(74%) | 11 ± 5<br>(85 %) | 6.2 ± 2.3<br>(83 %) | 4.2 ± 1.3<br>(96 %) |
| R69A | n.d. | > 1000<br>(35 %) | 31 ± 12<br>(60 %) | 20 ± 2.9<br>(88 %) |

The $K_d$ values were obtained by fitting the data to the quadratic equation. The average $K_d$ values with standard deviations were calculated from at least three independent experiments. When the fraction bound at the highest protein concentration was lower than 40%, the $K_d$ value was estimated as higher than 1 μM.
<sup>a</sup> – data from Table 1.
n.d. - binding was not detected up to 1 μM concentration of the protein

### K35 and R69 indirectly contribute to RNA binding at the central RNA binding site

We hypothesized that the differential effects of R69A and K35A on the binding of ProQ^NTD^ to distinct RNAs could result either from disruptions of direct contacts formed by K35 and R69 with specific RNAs or from indirect effects on alterations of contacts with other amino acids. Because R69 and K35 are located outside of the central concave-face pocket that has been proposed to interact with dsRNA (Pandey et al. 2020; Kim et al. 2022), we wondered whether R69 and K35 could form contacts with single-stranded regions on the 5ʹ and 3ʹ side of terminator hairpin, respectively. We reasoned that if K35 or R69 directly contacted a single-stranded region outside of the hairpin, a shortened RNA would show reduced affinity with WT ProQ, the mutant protein (K35A or R69A) would show reduced affinity with full-length RNA, but no additional loss of interaction would be observed when the mutant protein bound shortened RNA. To conduct these double-mutant experiments, we designed model RNA constructs derived from *cspE*-3ʹ and compared the effects of their 5ʹ and 3ʹ truncations on binding to WT ProQ^NTD^ and its R69A and K35A mutants (Fig. 6, Table 3, Suppl. Fig. S14). First, we constructed a model RNA (*cspE*-mini) by removing the minimally structured 17-nt from its 5ʹ end and introducing guanosines in three locations within this model construct to enable efficient *in vitro* transcription of further 5ʹ truncated constructs—at the 5ʹ end, the base of the terminator hairpin, and above the A-U base pairs at the base of the hairpin (Fig. 6A). The *cspE*-mini construct was found to bind WT ProQ^NTD^ with the same affinity as unmodified *cspE*-3ʹ (Fig. 6B, Table 3), suggesting that the 17-nt stretch of nucleotides on the 5ʹ end of *cspE*-3ʹ does not markedly contribute to RNA binding by WT ProQ^NTD^. Similarly, the loss of these 5ʹ nucleotides did not decrease the affinity of the R69A variant. Interestingly, however, the *cspE*-mini RNA bound to the K35A mutant with a ∼2-fold weakened binding affinity (Fig. 6B, Table 3, Suppl. Fig. S14), suggesting that loss of a longer 5ʹ single-stranded tail makes RNA binding more dependent on the K35 side chain. While there were only modest effects of removing the first 17 5ʹ nts, removal of the remaining single-stranded sequence on the 5ʹ side (Fig. 6A) had a dramatic effect on binding across the board: the resulting *cspE*-mini-5ʹ-blunt construct only achieved about 20% fraction bound with WT ProQ^NTD^ and resulted in no detectable binding to either R69A or K35A mutants (Fig. 6B, Table 3, Suppl. Fig. S14). To explore if releasing the 3ʹ-oligoU tail of *cspE-*3ʹ would improve binding to ProQ^NTD^ in the absence of nucleotides 5ʹ of the hairpin, we further removed four adenosines opposite the 3ʹ-oligoU tail (*cspE*-mini-5ʹ-truncated). Despite its longer single-stranded oligoU tail, no binding of this construct was detected even for WT ProQ^NTD^. Together, these data show that single-stranded nucleotides 5ʹ-adjacent to the terminator hairpin are essential for the RNA binding by ProQ^NTD^, while nucleotides further upstream of the terminator hairpin do not markedly affect RNA binding to WT ProQ but may make additional contacts to the protein that help compensate for the loss of K35. Importantly, these data are not consistent with either R69 or K35 directly contacting the 5ʹ single-stranded RNA.

**Figure 6.**
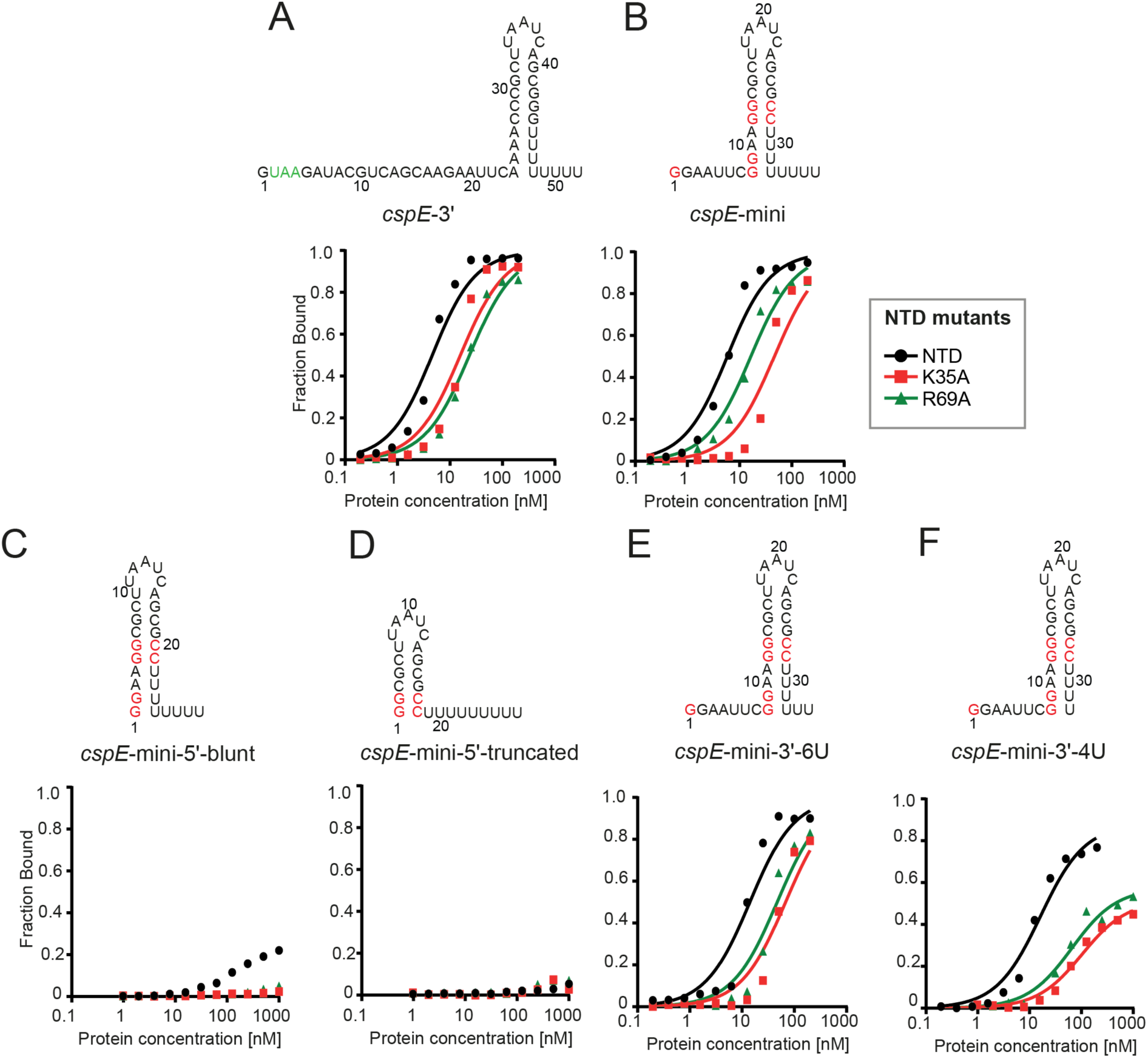
The substitutions of peripheral residues K35 and R69 have indirect effect on the RNA binding in the central pocket of the ProQ^NTD^. The RNA secondary structures were predicted using *RNAstructure* software. The data sets in which maximum fraction bound was above 40% were analyzed by fitting to the quadratic equation. Data for *cspE*-3ʹ are the same as in Figs. 3 and 4. The average equilibrium dissociation constant (*K*_d_) values are shown in the Table 3, and raw gel data are presented in the Suppl. Fig. S14.

**Table 3.** Equilibrium binding of *cspE*-3’-derived model RNAs to the ProQ NTD and its K35A and R69A mutants.

| ProQ<sup>NTD</sup><br>mutation | $K_d$ (nM)<br>(max. fraction bound %) | | | | | |
| --- | --- | --- | --- | --- | --- | --- |
|  | *cspE*-3' <sup>a</sup> | *cspE*-mini | *cspE*-mini-5'-blunt | *cspE*-mini-5'-truncated | *cspE*-mini-3'-6U | *cspE*-mini-3'-4U |
| wt NTD | 4.2 ± 1.3<br>(96 %) | 5.1 ± 1.1<br>(91 %) | >1000<br>(25 %) | n.d. | 13 ± 2.7<br>(91 %) | 16 ± 1.2<br>(81 %) |
| K35A | 18 ± 3.9<br>(88 %) | 44 ± 11<br>(86 %) | n.d. | n.d. | 63 ± 8.4<br>(82 %) | 75 ± 38<br>(41%) |
| R69A | 20 ± 2.9<br>(88 %) | 20 ± 7.7<br>(87 %) | n.d. | n.d. | 46 ± 8.4<br>(80 %) | 99 ± 35<br>(56 %) |

The $K_d$ values were obtained by fitting the data to the quadratic equation. The average $K_d$ values with standard deviations were calculated from at least three independent experiments. When the fraction bound at the highest protein concentration was lower than 40%, the $K_d$ value was estimated as higher than 1 μM.
<sup>a</sup> – data from Table 1.
n.d. - binding was not detected up to 1 μM concentration of the protein.

To explore whether R69 or K35 could affect RNA binding via contacts with the 3ʹ oligoU tail, we designed another construct (*cspE*-mini-3ʹ-6U), the 3ʹ tail of which was shortened from 8 to 6 Us, only two of which are predicted to be single-stranded (Fig. 6C). This 3ʹ truncation weakened the binding of WT ProQ^NTD^ an additional 2.5-fold beyond *cspE*-mini (Table 3), confirming previous observations that the length of the 3ʹ U-tail is important for the RNA binding to the FinO domain (Stein et al. 2020). The 3ʹ truncation also further reduced the binding affinity of both R69A and K35A mutants (by ∼2.4-and ∼1.4-fold), suggesting that neither of these residues primarily functions by contacting the terminal portion of the 3ʹ U tail.

We next examined the effects of a more extensive 3ʹ truncation, removing an additional 2 terminal uridines to create *cspE*-mini-3ʹ-4U (Fig. 6C). Binding of all three ProQ^NTD^ variants (WT, R69A and K35A) was further weakened to this RNA construct, with strong effects seen on the maximum fraction bound for R69A and K35A proteins (Table 3). The fact that substitution of either R69 or K35 had additional detrimental effects on the binding of *cspE*-mini-3ʹ-4U beyond those of *cspE*-mini-3ʹ-6U suggests that these two nucleotides do not directly contact K35 or R69. We note that the effect of the 4-uridine truncation in *cspE*-mini-3ʹ-4U is smaller than previously observed for a different *cspE*-3ʹ construct with the same truncation (Stein et al. 2020). While the affinities of these two constructs indeed differ in side-by-side binding assays (Suppl. Fig. S15), the result that a 4-uridine truncation is detrimental for RNA binding is consistent across both constructs.

In summary, these mutational studies suggest that R69 and K35 both likely contribute to contacts to the core terminator hairpin of the RNA rather than the adjacent 5ʹ or 3ʹ sequences. This is consistent with the above observation that the heightened susceptibility of SibB to R69A and K35A mutations is explained by the properties of its terminator hairpin (Fig. 5, Table 2). In addition, K35 may contribute in part to contacts that become especially important when RNAs possess a shorter extension on the 5ʹ side of their terminator hairpin.

## DISCUSSION

In this study, we set out to refine our model of molecular recognition of RNA ligands by the FinO-domain of *E. coli* ProQ. Data collected from both biochemical and genetic experiments support a model in which all essential RNA-binding residues are located on the concave face of ProQ’s N-terminal FinO domain. This includes basic and aromatic side chains poised in the concave-face pocket to interact with intrinsic terminators at the 3ʹ end of many of ProQ’s RNA ligands. Our data suggest that additional residues on the periphery of the concave face—and even pointing towards the opposing convex face of the FinO domain—make smaller contributions to RNA binding that are more varied across RNA ligands. Together, these findings strengthen and extend our prior results, showing that nearly all the ProQ residues implicated previously in RNA binding through *in vivo* methods indeed impact RNA binding directly *in vitro*, and allowing for nuanced comparisons to be drawn across a variety of RNA ligands.

### Revisiting the structure of *E coli* ProQ’s FinO Domain

One of the questions at the outset of this work was how to explain that amino acids required for RNA binding in *E.coli* ProQ appeared to be quite distant from one another on the available structural model of the protein; specifically, a universally conserved arginine (R80 in *Ec* ProQ) was the lone RNA-binding residue predicted to fall on the convex face of the *E. coli* protein. Because available structural models positioned this residue differently (Gonzalez et al. 2017; Jumper et al. 2021; Varadi et al. 2022), we elected to take an unbiased approach to elucidate the *in vivo* position of R80 in *Ec* ProQ. A compensatory mutagenesis screen identified several amino acid substitutions that partially rescue the detrimental effects of an R80K substitution. The position of these compensatory substitutions are most consistent with a structural model in which the more conserved face of the FinO domain of *Ec* ProQ possesses a concave pocket containing both R80 and Y70, analogous to the concave-face pocket seen in other structural homologs (Suppl. Fig. S2) as well as in the AlphaFold model for *Ec* ProQ. For this reason, we used the structure of *E. coli* ProQ predicted by AlphaFold for downstream structural modeling in this work (Jumper et al. 2021; Varadi et al. 2022). There are several possibilities for why the position of R80 in *E. coli* ProQ could be different inside the cell and in the AlphaFold model than it was in the initially reported NMR structure (Gonzalez et al. 2017). For instance, it is possible that multiple protein conformations are accessible in the absence of RNA, but that the conformation with a concave-facing R80 residue is locked in place once the protein binds to RNA. Alternatively, conformational flexibility in the region of the protein containing R80 could make its position difficult to resolve by NMR.

While the most important takeaway from the compensatory substitutions isolated in our forward genetic approach was to examine the proximity of the mutated residues to the position of R80 in each of the available structural models (Fig. 2D, E), it is also interesting to consider what the specific substitutions reveal about the structure of ProQ in this region. To begin with, the AlphaFold model provides an explanation for why the seemingly conservative R80K substitution is so strongly deleterious to RNA binding *in vivo* and *in vitro* (Figs. 1B, C and 3). While both lysine and arginine are positively charged at physiological pH, the aliphatic portion of lysine’s side chain is shorter than that of arginine. If the side chain of R80 points through to the concave face from the β-hairpin containing the residue (Fig. 2E), substitution with lysine could impact RNA binding either directly by moving the residue’s charged terminus away from the RNA, or indirectly by destabilizing the hydrophobic core of the protein. Several of the validated compensatory substitutions (V74L, L91V and L91P) could subtly tweak the hydrophobic environment surrounding R80 to create space for, or help to reposition, lysine’s primary amine. Two additional substitutions at V74 could potentially stabilize lysine’s primary amine – through either a cation-π interaction (V74F) or a hydrogen bond (V74K, Fig. 2E). It is also possible that the presence of two neighboring primary amines in the R80K-V74K double mutant recreates some of the “softer” character of R80’s guanidinium group. Importantly, even the strongest of the identified compensatory substitutions (L91V and V74K) lead to only modest recovery of RNA binding by the R80K ProQ mutant. This underscores the key role that R80 plays in FinO-domain structures, consistent with its universal conservation.

### Structural Model of *E. coli* ProQ-RNA Interactions

As noted above, the AlphaFold structural model for *E. coli* ProQ shares many structural features with the other four FinO-domain homologs with solved structures (Suppl. Fig S2) (Ghetu et al. 2000; Chaulk et al. 2010; Immer et al. 2020; Kim et al. 2022). Only one structure solved to date has captured high-resolution details of the FinO domain in complex with an RNA ligand: the recent crystal structure of *L. pneumophila* RocC bound to the terminator hairpin of the RocR RNA (Kim et al. 2022). This structure, which shows RocC residues making multiple contacts with the 3ʹ side of a dsRNA duplex and a 3ʹ single-stranded tail, offers several important insights for the interpretation of the genetic and biochemical data presented here. The AlphaFold model of *Ec* ProQ^NTD^ aligns very well with the structure of RocC (RMSD = 1.09 Å; Fig. 7A), providing a possible structural model for *Ec* ProQ interacting with a terminator hairpin (Fig. 7B). This alignment-based model is consistent with the residues we have identified as most important for RNA binding: K54, R58, R62, Y70 and R80 are all found in the immediate vicinity of the terminator hairpin (Fig. 7B). Figures 7C and 7D summarize the effects of substitutions on the concave face of *Ec* ProQ^NTD^, comparing results from *in vitro* and *in* vivo binding assays (Figs. 3, 4; Pandey 2020). Overall, there is good alignment between *in vivo* and *in vitro* results, suggesting that most residues identified through forward and reverse genetics exert their effects on RNA binding directly, rather than through downstream effects like recruitment of another protein or complex.

**Figure 7.**
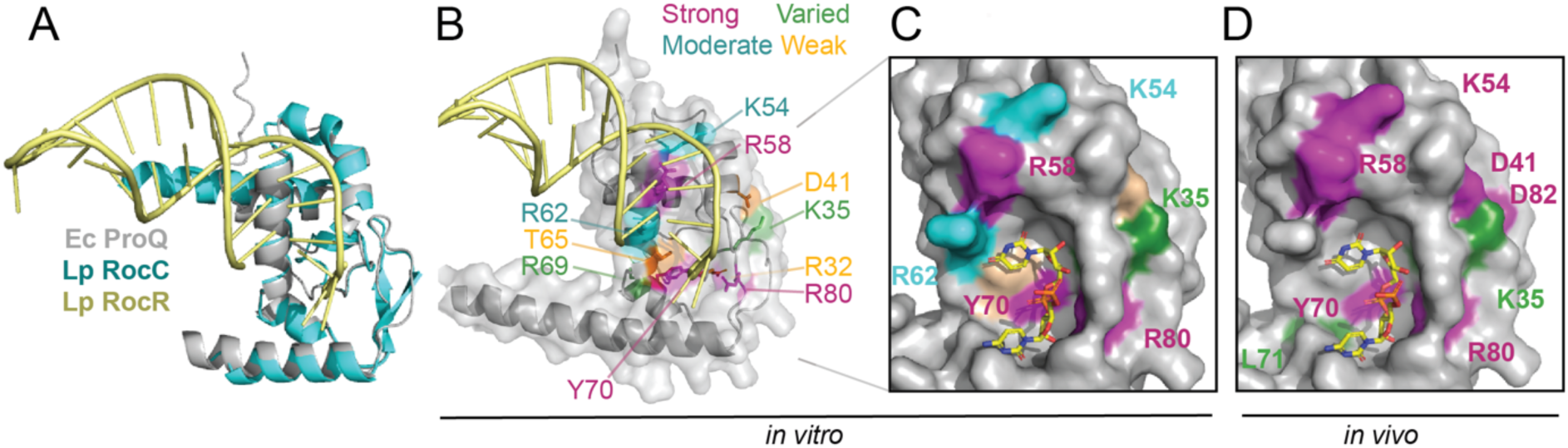
Model for ProQ-RNA interaction. (A) Alignment of AlphaFold model of FinO domain of ProQ (Jumper et al. 2021; Varadi et al. 2022) and *Lp* RocC/RocR co-crystal structure (PDB ID: 7RGU; (Kim et al. 2022)). (B) Summary of *in vitro* biochemical results. Residues which were modified for gel-shift experiments are colored based on the size of the detrimental effects on RNA binding when modified: strong (R80, Y70, R58; magenta), moderate (K35, K54, R62; cyan) and effects that varied across RNA ligands (K35, R69; green). Not labeled: residues whose substitution had only weak effects (D41, T65). (C, D) Surface representation of the concave-face binding pocket of ProQ, with residues colored as in (B). The terminal two pyrimidine nucleotides from RocR are shown in stick representation and are colored by atomic identity. The two panels compare (C) ProQ residues identified as important for RNA interactions *in vitro* in the current work and (D) residues found in previous B3H experiments to contribute to *cspE*-3’ and SibB binding *in vivo* (Pandey et al. 2020).

In the X-ray structure of *Lp* RocC, the 3ʹ terminus of the RocR RNA in is bound by residues corresponding to *E. coli* amino acids Y70 and R80, as well as the polypeptide backbone surrounding G37 (Kim et al. 2022). Each of these residues directly contacts the RNA: both the arginine and tyrosine make hydrogen bonds to the final internal phosphate moiety, the arginine makes an additional hydrogen bond with the 2ʹ OH of the penultimate nucleotide, and the backbone amides surrounding glycine contact the terminal 3ʹ-OH group. The fact that the shape of the concave pocket is almost identical between *Lp* RocC and *Ec* ProQ suggests that *Ec* ProQ may also bind the 3ʹ terminal two nucleotides in this pocket in the vicinity of Y70 and R80 (Fig. 7C, D).

This tyrosine is one of the most highly conserved residues in FinO-domain structures and has been previously implicated as critical for RNA binding (Attaiech et al. 2016; Immer et al. 2020; Pandey et al. 2020; El Mouali et al. 2021b). Even a conservative substitution from tyrosine to phenylalanine eliminates RNA binding *in vitro* (Fig. 3, Table 1) and *in vivo* (Pandey et al. 2020), underscoring the essential role of this hydroxyl group on the concave face. Finally, both neighboring amino acids—R69 and L71—have been found to contribute to RNA binding in ProQ or its homologues (Fig. 4, Table 1) (Attaiech et al. 2016; Pandey et al. 2020; El Mouali et al. 2021b; Rizvanovic et al. 2021), which suggests that these residues could influence RNA binding through effects on the position of Y70.

The other universally conserved amino acid residue that binds the 3ʹ terminus of the RNA in the RocC/R co-crystal structure is *E. coli* R80. Our work shows that substitution of this arginine with either lysine or alanine severely impairs RNA binding *in vitro* (Fig. 3, Table 1) and *in vivo* (Fig. 1; Pandey et al. 2020)). The homologous arginine residue in *Lp* Lpp1663 underwent chemical shift changes in NMR experiments in the presence of RaiZ or oligoU RNA (Immer et al. 2020; Kim et al. 2022). There is also considerable genetic evidence for the importance of this residue: multiple substitutions at this position (serine, histidine and cysteine) were identified in *Ec* ProQ in an unbiased B3H genetic screen (Pandey et al. 2020) and, indeed, no other residue was found to restore any detectable interaction at this position (Suppl. Table S1). In addition, two independent genetic screens looking for mutations abrogating the role of ProQ in gene expression regulation in *S. enterica* isolated substitutions of R80 to histidine, serine or glycine (El Mouali et al. 2021b) or to histidine (Rizvanovic et al. 2021).

The importance of the precise shape of the concave pocket containing Y70 and R80 in *Ec* ProQ is supported by the prior identification of the deleterious alanine substitution of the universally conserved G37 in our *in vivo* forward genetic screen (Pandey et al. 2020), suggesting that flexibility in the polypeptide backbone is required in this region. Interestingly, G37 neighbors the K35 residue, which makes varying contributions to binding across different RNAs (Fig. 4, Table 1). It is therefore possible that mutation of K35 indirectly affects RNA binding by influencing the molecular environment of the G37 residue within this RNA-binding pocket.

The double-stranded stem of the RNA terminator hairpin is contacted in the *Lp* RocC co-crystal structure by residues within the H3 helix of the FinO domain (Kim et al. 2022). Alignment of the *Ec* ProQ AlphaFold model (Fig. 7A) places the terminator hairpin’s major groove and phosphate backbone in the vicinity of *Ec* H3 residues K54, R58 and R62, each of which makes moderate or strong contributions to RNA binding *in vitro* (Figs. 3, 7B, C). Of these H3 residues, we found substitutions at R58 to be especially damaging to RNA interactions (Fig. 4, Table 1). In agreement with the importance of this residue, the R58A mutation in *Ec* ProQ also had a strong negative effect on RNA binding in the B3H assay (Pandey et al. 2020) and a cysteine introduced to the equivalent position in the FinO protein crosslinked with FinP RNA (Ghetu et al. 2000). The other H3 residues examined here (K54 and R62) are both located in evolutionarily variable positions (Suppl. Fig. S1) and make moderate contributions to RNA binding *in vitro* (Fig. 4, Table 1). Notably, a lysine in the position in *Lp* RocC corresponding to *Ec* K54 is part of the proposed N-cap motif that recognizes the double-stranded portion of the RNA hairpin by contacting phosphate groups (Kim et al. 2022). While an alanine substitution of K54 in *Ec* ProQ was found to be detrimental to RNA binding in the B3H assay (Pandey et al. 2020), this position was not identified in any of the forward genetic screens conducted for *Ec* ProQ or homologous proteins. Like K54, no substitutions at R62 were identified in any of the previously conducted genetic screens, but a cysteine introduced to the corresponding position of the FinO protein crosslinked to RNA (Ghetu et al. 2002). Overall, the fact that substitution of non-conserved residues such as K54 and R62 still produces moderately detrimental effects on RNA binding *in vitro* suggests that these residues could play a role in RNA contacts that are specific to a subset of FinO-domain proteins, including *Ec* ProQ. R62 is particularly interesting given that it is not conserved in *Lp* RocC, but its position in the alignment-based model (Fig. 7B) suggests it may be able to interact with a backbone or base in the 5ʹ side of the terminator hairpin RNA. Such contacts to the 5ʹ side of the hairpin are notably missing from the *Lp* RocC/R structure and further structural studies with the *E. coli* ProQ protein will be necessary to determine if R62 indeed plays such a role.

The weak effects of substitutions of some conserved amino acids on RNA binding observed here are also consistent with a lack of direct RNA contacts seen in the RocC/RocR co-crystal structure (Fig. 4, Table 1) (Kim et al. 2022). For instance, while R32 is located in a position that contains either arginines or lysines in homologous proteins (Suppl. Fig. S1), its side chain is found on the convex face, explaining why an alanine substitution did not markedly affect the RNA binding (Fig. 4, Table 1). On the other hand, T65 is located on the concave face, immediately adjacent in the structure to universally conserved Y70 in a position where homologous proteins contain residues with hydrogen-bonding potential (threonine, serine or cysteine). Despite this, alanine substitution of T65 did not affect RNA binding with purified components (Fig. 4, Table 1). This is consistent with the lack of effect on RNA binding of the corresponding threonine residue in *Lp* RocC (Kim et al. 2022). Another residue found to make only small contributions to RNA binding is D41. Alanine substitution of this residue was detrimental to RNA binding in our B3H assay (Pandey et al. 2020) and substitution of the corresponding D56 with tyrosine was detrimental for function in *Lp* RocC (Attaiech et al. 2016). However, a D41A mutation showed only weak effects on RNA binding in the purified system in this study (Fig. 4, Table 1), consistent with a lack of RNA contacts of the corresponding aspartate in the RocC/RocR structure (Kim et al. 2022). This difference between *in vivo* and *in vitro* effects could suggest that D41 plays an important *in vivo* role that is not reflected in binding assays with purified components, such as fine-tuning the positioning of RNA on the FinO domain and/or preventing interactions with other competing RNAs.

While several residues on the concave face of *Ec* ProQ were previously found to be important for the binding of two RNAs (the 3ʹ-UTR of *cspE* and the sRNA SibB; Pandey 2020), the data presented here furthers these observations by examining the effects of substitutions in a purified *in vitro* system and against a larger set of seven RNAs (Figs. 3, 4, Table 1, Suppl. Fig. S5). The fact that similar conclusions were reached for this larger set of RNAs suggests that many of the RNA ligands of *Ec* ProQ share a similar mode of binding as was seen in the prior genetic study (Pandey 2020). Indeed, most residues examined in this work had similar effects on all seven RNAs tested—whether strong (R58, Y70 and R80), moderate (K54, R62) or weak (R32, D41, and T65) (Figs. 3, 4, Table 1), suggesting that these residues likely recognize core RNA features that all seven RNA ligands have in common.

In contrast to most residues examined, substitutions of K35 and R69 stand out as having especially varied effects on RNA binding (Figs. 3, 4, Table 1, Suppl. Figs. S6-S12). K35 and R69 are positioned on the concave face and the edge of the convex face, respectively, and occupy evolutionarily variable positions within stretches of highly conserved sequence (Suppl. Figs. S1, S2). The differential contribution of *Ec* K35 to interaction with *cspE*-3ʹ and SibB has been previously observed in the B3H assay (Pandey et al 2020), but contributions of this residue have yet to be examined in other homologs. Likewise, the position of *Ec* R69 had not been previously implicated in RNA binding by other FinO-domain proteins. The amino acids found in this position differ widely across FinO-domain proteins and include basic, neutral and acidic residues (Suppl. Fig. 1). Given this diversity and the fact that the side chain of R69 points towards the convex face of *E. coli* ProQ, it is quite interesting that its substitution so strongly affects binding of some of the RNAs in our panel. Given the proximity of R69 to Y70, one possibility is that R69 exerts some of its differential effects on RNA binding through fine-tuning of the environment of Y70, analogous to a potential role of K35 acting through other residues within the RNA-binding pocket.

Across all ProQ^NTD^ mutants examined, *cspE*-3ʹ was consistently one of the strongest-binding RNAs and SibB one of the weakest (Figs. 3, 4, Table 1). This *in vitro* difference in binding strength is consistent with the previous *in vivo* observation that a K35A mutation in *Ec* ProQ^ΔCTD^ had a strong effect on the binding of SibB in the B3H assay, but only weakly affected binding of the *cspE* 3ʹ-UTR (Pandey et al. 2020). Tighter interaction of *cspE*-3ʹ RNA than SibB is also consistent with transcriptome-wide datasets probing *in vivo* interactions with WT ProQ^NTD^. For instance, *cspE*-3ʹ was consistently among the top 15 RNAs with 3ʹ terminators identified as ProQ ligands in *E. coli* using either RIL-seq or CLIP-seq and in *S. enterica* using CLIP-seq (Holmqvist et al. 2018; Melamed et al. 2020; Stein et al. 2020). On the other hand, SibB fell outside of the top 30 RNAs identified by CLIP-seq in *E. coli* and was not present among the top 50 RNAs in datasets from *E. coli* RIL-seq or *S. enterica* CLIP-seq experiments (Holmqvist et al. 2018; Melamed et al. 2020; Stein et al. 2020).

The differences in ProQ^NTD^ binding observed *in vitro* for RNAs beyond *cspE*-3ʹ and SibB have also been hinted at by previous studies. Following *cspE*-3ʹ and *cspE*81-3ʹ, the next strongest ProQ^NTD^-interacting RNA was *malM*-3ʹ, followed by SibA, *gapA*-3ʹ, RybB and finally SibB (discussed above). While *malM*-3ʹ was among the top-50 ProQ RNA ligands in two of the three datasets discussed above (*E. coli* RIL-seq and CLIP-seq), *gapA*-3ʹ and RybB were each identified in only one experiment (*E. coli* CLIP-seq or RIL-seq, respectively) (Holmqvist et al. 2018; Melamed et al. 2020; Stein et al. 2020). On the other hand, SibA was not among the top 50 ligands in any of the three datasets, though its antisense mRNA (*ibsA*) was present in the *E. coli* RIL-seq dataset (Melamed et al. 2020). Although identification in global profiling data may reflect other aspects besides binding affinity, including relative concentrations of RNAs, the data from global profiling studies (Holmqvist et al. 2018; Melamed et al. 2020) are also consistent with the conclusion that there are differences among RNAs in how well they are bound by ProQ.

Given the differences in binding to K35A and R69A mutants of ProQ^NTD^ observed across the seven RNAs, it is interesting to consider what features of the RNA ligands may drive these differences. A previous study showed that optimal binding of *malM*-3ʹ and *cspE*-3ʹ requires a terminal hairpin stem with single-stranded tail of at least four uridines (Stein et al. 2020). While all seven RNAs studied here have a terminator hairpin structure with a 3ʹ-oligoU tail and an A-rich motif on the 5ʹ side of the hairpin that could partially interact with the U-tail, the length and sequence of these elements differ across the seven RNAs (Suppl. Fig. S5). The structure of the terminator hairpin itself appears to modulate the sensitivity RNA binding to substitutions on the periphery of the FinO-domain: replacing the bottom part of the terminator hairpin and 3ʹ tail of SibB with that from *cspE*-3ʹ strongly improved its binding to R69A mutants (Fig. 5, Table 2, Suppl. Fig. S13). The diversity of sequences around the terminator hairpin further contributes to the strengths of RNA interaction with ProQ^NTD^ and their sensitivities to substitutions of K35 and R69. For instance, replacing the two terminal cytidines of SibB with uridines markedly improved the binding of the resulting RNA molecule (Fig. 5, Table 2), while shortening the region on either the 5ʹ or 3ʹ side of the terminator within a minimal *cspE*-3ʹ RNA weakened its interaction with ProQ (Fig. 6, Table 3, Suppl. Fig. S14). RNA structure upstream of the terminator hairpin may also modulate the strength of RNA binding. A shorter variant of *cspE*-3ʹ was less affected by K35A and R69A mutations than a longer variant with a stable structure on the 5ʹ side of its terminator hairpin (Fig 6, Table 3), suggesting that adjacent RNA structure may restrict the terminator’s positioning on ProQ^NTD^.

Despite the sensitivity of RNAs to K35A and R69A substitutions being dictated by sequences within and around their terminator hairpins, the analysis of either mutant protein binding to minimal *cspE*-3ʹ RNAs was not consistent with K35 or R69 directly contacting RNA on either side of the terminator hairpin (Fig. 6, Table 3). Rather, it is more likely that these residues exert their effects on RNA interaction by modulating the way the primary RNA-binding site (*e.g.* Y70 and R80) contacts the base of the terminator stem and the 3ʹ U-tail. The fact that the seven RNAs studied here show a similar rank-order in their interaction strengths to WT, K35A and R69A ProQ^NTD^ is consistent with the absence of K35 and R69 amplifying inherent differences in the ability of different RNAs to bind ProQ^NTD^. In summary, our data suggest that an RNA’s terminator hairpin structure and the nucleotides surrounding it dictates the strength of that RNAs interaction with ProQ^NTD^ and its susceptibility to mutations.

### Outlook

The work presented here significantly extends our understanding of how the concave face of the FinO-domain of *E. coli* ProQ recognizes RNA ligands. Our work supports a model in which the structure near the base of an RNA’s terminator hairpin dictates the strength of its interaction with ProQ through interactions of the dsRNA with K54, R58 and R62 and its uridine tail with the pocket formed between Y70 and R80. Even with this improved understanding of interactions between RNA terminators and the FinO domain of *E. coli* ProQ, there are many important questions about this class of RNA-binding proteins to be investigated in future work. For instance, different FinO-domain proteins are known to show different patterns of RNA interaction *in vivo* from CLIP-seq and other transcriptome-wide methods (Attaiech et al. 2016; Melamed et al. 2016; Smirnov et al. 2016; Holmqvist et al. 2018; Bauriedl et al. 2020; Gerovac et al. 2020; El Mouali et al. 2021a). It will be important to explore what modifications are present in the RNA-interacting residues of these homologs to mechanistically enforce the observed differences in RNA-binding patterns. In addition, this work has focused exclusively on the interaction of *E. coli* ProQ’s N-terminal FinO domain with RNAs that include intrinsic-terminator-like 3ʹ ends. There are interesting unanswered questions about how ProQ may interact with 5ʹ UTRs, as has been observed *in vivo* (Holmqvist et al. 2018; Melamed et al. 2020). It will also be important to determine the role that the unstructured linker and C-terminal Tudor domain may play in the function of *E. coli* ProQ. While these domains are not critical for RNA binding *in vivo* or *in vitro*, there is intriguing genetic evidence in *Salmonella* that the CTD may play a critical role in a function of ProQ downstream of RNA binding (El Mouali et al. 2021b; Rizvanovic et al. 2021). The CTD or linker could perhaps play roles in facilitating RNA-RNA interactions between RNAs bound to the N-terminal FinO domain or in recruiting other factors within the cell. Our hope is that the improved mechanistic understanding of how the FinO-domain of *E. coli* ProQ interacts with RNA terminators will facilitate future work to probe additional unanswered questions about the molecular mechanisms and gene-regulatory behavior of *E. coli* ProQ and the broader class of FinO-domain proteins.

## MATERIALS AND METHODS

### Bacterial 3-hybrid (B3H) assay

*E. coli* strains, plasmids and oligonucleotides used in the B3H assay are listed in Suppl. Tables S2-S4. NEB5α, purchased from New England Biolabs, was the recipient strain for all cloned B3H plasmids. KB473 served as the reporter strain for all β-galactosidase (β-gal) assays. Each strain and plasmid has specific antibiotic resistance gene(s), listed with the following abbreviations: AmpR (ampicillin and carbenicillin), CmR (chloramphenicol), KanR (kanamycin), StrR (streptomycin), and TetR (tetracycline). All strains were stored as glycerol stocks at –80°C.

In this B3H system, plasmids express three hybrid components: 1) a DNA-RNA adapter protein, CI-MS2^CP^, tethers 2) a Bait RNA construct upstream of a test promoter such that it is available for interaction with 3) an RNA Polymerase (RNAP)-tethered prey protein (Fig 1A). Reporter cells encode a *lacZ* reporter gene downstream of a test promoter on a single-copy Fʹ episome. Transformation of reporter cells with all three plasmids (pPrey, pAdapter, and pBait) leads to a boost in β-gal levels relative to basal levels indicated by three negative controls in which half of each hybrid component is left out (Fig 1B); the strength of an RNA-protein interaction correlates to the fold-stimulation in β-gal activity over basal levels when all components are present (Fig 1C; (Wang et al. 2021; Stockert et al. 2022)).

### β-galactosidase assays

Liquid assays were performed with the use of the bacterial three-hybrid system. Reporter cells (KB473) were co-transformed with pAC-, pBR-, and pCDF-derived plasmids. pAC constructs express the CI-MS2^CP^ fusion protein, while pCDF-pBAD constructs express the MS2^hp^ fusion RNA and pBR-α expresses the α-ProQ fusion protein. For each transformation, there were three controls, one where each of these core plasmids was replaced with an “empty” construct. Single colonies from each transformation were inoculated into 1 mL of LB broth supplemented with 0.2% arabinose and antibiotics: carbenicillin (100 μg/mL), chloramphenicol (25 μg/mL), kanamycin (50 μg/mL), and spectinomycin (100 μg/mL) in a 2 mL 96 well deep-well block (VWR) sealed with breathable film (VWR) and shaken at 900 rpm and 37°C overnight. Overnight cultures were back-diluted (1:40) into 200 μL LB supplemented with the same antibiotics and arabinose as outlined above, as well as 0 μM, 5 μM, or 50 μM IPTG (isopropyl-ß-D-thiogalactoside; see figure legends) into optically clear 200 μL flat-bottom 96-well plates covered with plastic lids (Olympus). Mid-log cells (OD_600_ 0.3-0.6) were transferred into a new 96-well plate with rLysozyme and PopCulture reagent (EMD Millipore) and allowed to lyse for 0.5-4 hours. Lysate was transferred into a fresh optically clear 96 well plate (Olympus) with Z-buffer, ONPG (O-nitrophenyl-ß-D-galactopyranoside), and ß-mercaptoethanol. β-gal activity was measured by taking OD_420_ values every minute at 28°C for 1 hour using a microplate spectrophotometer (Molecular Devices SpectraMax). OD_420_ readings were normalized using the OD_600_ values from directly before lysis to give β-gal activity in Miller units (Thibodeau et al. 2004; Stockert et al. 2022). β-gal activity was averaged over three replicates for each experimental condition and then divided by the highest relevant negative control to give the fold-stimulation. Negative controls refer to transformations in which one component of the assay (pPrey, pBait, or pAdapter) has been replaced with an empty construct. Error for fold was propagated from the standard deviations of experimental and negative control averages. Assays were conducted in biological triplicate on at least three separate days. For qualitative plate-based assays, 3.2 μL of mid-log cells from above were pipetted from the 96-well plate onto a large LB agar plate supplemented with inducers (0.2% arabinose and 1.5 μM IPTG), antibiotics (carbenicillin (100 μg/ml), chloramphenicol (25 μg/ml), kanamycin (50 μg/ml) and spectinomycin (100 μg/ml)) and indicators (X-gal (40 μg/ml) and TPEG (200 μM)). Plates were incubated at 37°C overnight and then moved to a 4°C fridge for at least one day before pictures were taken of the plate.

### Western Blots

Cell lysates from β-gal assays were normalized based on OD600 with LB plus PopCulture Reagent. Lysates were mixed with 6× Laemmli loading dye, boiled for 10 min at 95°C and electrophoresed on 10–20% Tris–glycine gels (Thermo Fisher) in 1× NuPAGE MES Running Buffer (Thermo Fisher). Proteins were transferred to PVDF membranes (BioRad) using a semi-dry transfer system (BioRad Trans-blot Semi-Dry and Turbo Transfer System) according to manufacturer’s instructions. Membranes were probed with 1:10000 primary antibody anti-ProQ overnight at 4°C and then a horseradish peroxidase (HRP)-conjugated secondary antibody (anti-rabbit IgG; 1:10000). Chemiluminescent signal was detected using ECL Plus western blot detection reagents (Bio-Rad) and a c600 imaging system (Azure) according to manufacturer’s instructions

### Library Construction for Saturation Mutagenesis

Single site mutagenesis was performed using Q5 site-directed mutagenesis. Mutagenic forward and reverse primers were designed using NEBaseChanger. For the Y70X and R80X library plasmids, a region of the primer representing the single codon of interest was replaced by a mixture of 25% of each nucleobase in each of the three sites, written as “NNN” in the primer sequence (Suppl. Table S4). Otherwise, the primers were complementary to the backbone sequence. For multi-site saturation mutagenesis of the R80K vector, multiple forward primers were designed to mutagenize one codon position at a time with a degenerate NNN codon (Suppl. Fig. S4A). For each of three libraries targeting a different 6-7aa region of ProQ, 6-7 forward primers were used in a pooled PCR reaction with a single reverse primer. Primers were used in a polymerase chain reaction (PCR) with 2X Phusion Master Mix (New England Biolabs) and the appropriate parent plasmid: pKB955 (pBrα-ProQ^ΔCTD^) for Y70X and R80X libraries and pSP144 (pBrα-ProQ^ΔCTD^-R80K) for the R80K multisite compensatory libraries. For Y70X and R80X libraries, PCR products were Kinase-Ligase-DpnI (KLD) treated in 10μL reactions containing 1 μL PCR Product, 1 μL T4 DNA Ligase (New England Biolabs), 1 μL T4 DNA Ligase buffer (New England Biolabs), 1 μL T4 polynucleotide kinase (PNK; New England Biolabs), 1 μL DpnI (New England Biolabs), and 5 μL MilliQ water. For the multisite-ProQ-R80K libraries, commercial KLD mixture was used according to manufacturer’s instructions (New England Biolabs). KLD products for each library were transformed into NEB5α cells. Cells were serially diluted and spread on LB-carbenicillin and incubated at 37°C overnight to provide a near-lawn for library preparation and to allow for estimates of colony numbers (Suppl. Table S5). Plasmid for each library was directly miniprepped from a cell slurry harvested from the overnight plates and stored at –20°C for further use.

### B3H Screening

Screens of these library plasmids were performed with the use of the B3H assay. The plasmid libraries were transformed into eKB473 reporter cells pre-transformed pKB1210 (pBait-*malM*-3ʹUTR) and pCW17 (pAdapter). Cells were heat-shocked and plated on LB agar supplemented with inducers (0.2% arabinose and 1.5 μM IPTG), antibiotics (carbenicillin (100 μg/ml), chloramphenicol (25 μg/ml), kanamycin (50 μg/ml) and spectinomycin (100 μg/ml)) and indicators (X-gal (40 μg/ml) and TPEG (200 μM)). Plates were incubated at 37°C overnight. Each time that a transformation was performed, positive and negative controls were transformed alongside the experimental conditions to allow for comparison of blue/white levels (positive: wild-type ProQ; negative: alpha empty, pBrα or pSP144, ProQ-R80K). The plates were transferred from the 37°C incubator to 4°C once colonies had grown to sufficient size, ∼18 hours. Plates were stored at 4°C for a minimum of ∼5 hours and a maximum of ∼3 days before being examined for the presence of blue colonies. Colonies that showed any blue color above that of pPrey-empty negative-control colonies were restruck on indicator plates to ensure single blue colonies. Plasmid from these colonies were miniprepped and sent for sequencing. Sequences were aligned to the sequence of wild-type *E. coli ProQ* to identify mutations at the codon of interest.

### Preparation of overexpression constructs and protein purification

The sequences of *E. coli* N-terminal domain (NTD; residues 1-130) of ProQ were cloned into pET15b vector (Novagen), as described (Stein et al. 2020). In the construct the coding sequence of the protein was preceded by His_6_-tag and TEV protease recognition sequence (ENLYFQ↓S), so an additional serine residue remains after the cleavage of the N-terminal His_6_-tag. The mutated variants of NTD of ProQ were obtained by site-directed mutagenesis, where the substitutions were introduced by specifically designed primers (Suppl. Tables S6, S7) to change a single amino acid in the protein sequence. All constructs were overexpressed in BL21 Δ*hfq* strain (a kind gift of Prof. Agnieszka Szalewska-Pałasz, University of Gdańsk), and purified as described (Stein et al. 2020). The samples were stored in buffer (50 mM Tris, pH 7.5, 300 mM NaCl, 10% glycerol and 1 mM EDTA) at −80°C as 10 and 20 μl aliquots and used without refreezing. The concentration of proteins was determined by measuring the absorption at 280 nm using extinction coefficient of 9650 M^-1^ cm^-1^.

### RNA preparation

The DNA templates used for *in vitro* transcription were obtained by Taq polymerase extension of chemically synthesized overlapping oligodeoxyribonucleotides (Sigma-Aldrich, Metabion, Suppl. Table S8). RNA molecules were transcribed with T7 RNA polymerase and purified using denaturing gel electrophoresis, as described (Milligan et al. 1987; Olejniczak 2011). In the next step RNAs were 5’-^32^P labeled using T4 polynucleotide kinase (Thermo Scientific), followed by phenol-chloroform extraction, purification on denaturing gel and precipitation with ethanol.

### In vitro RNA binding (gel shift) assay

Prior to use, RNA molecules were denatured for 2 min in 90°C followed by 5 min refolding on ice. The concentration series of all proteins were prepared by 2-fold dilutions from the highest concentration (given above the gel image). In each binding reaction, 1 nM ^32^P-labeled RNA was mixed with a protein sample diluted in binding buffer (25 nM Tris, pH 7.5, 150 nM NaCl, 5% glycerol, 1 nM MgCl_2_), and incubated for 30 min at RT in low-protein binding microplates pre-treated with a solution containing 0.0025% bovine serum albumin (BSA). After this time, reactions were loaded onto a 6% polyacrylamide gel (19:1) running in 0.5x TBE buffer at 4°C. After the electrophoresis, gels were vacuum-dried and exposed to phosphor screens overnight. The signal was quantified using a phosphorimager and MultiGauge software (Fuji FLA-5000), data were fitted to a quadratic equation using GraphPad Prism software and the equilibrium dissociation constant (*K*_d_) values were calculated as described (Stein et al. 2020). The average *K*_d_ values given in tables were calculated based on at least three independent experiments.

## Supplemental material

Supplemental material is available for this article.

## Author contributions

E.M.S. analyzed RNA binding by ProQ^NTD^ *in vitro*, and cloned and purified all mutant proteins; Su.W. and K. D. conducted B3H screens; Su.W. and C.M.G. analyzed RNA binding using the B3H assay; Su.W., K. D, Sh. W., and K.E.B. analyzed FinO domain structures; M.O. and K.E.B. analyzed data and wrote the paper.

## ACKNOWLEDGEMENTS

We thank Gisela Storz for helpful discussions about this manuscript. This work in the M.O. lab was supported by National Science Centre in Poland [grant No. 2018/31/B/NZ1/02612]. This work in the K.E.B lab was supported by National Institutes of Health [R15GM135878], the Henry R. Luce foundation, the Camille and Henry Dreyfus Foundation and Mount Holyoke College. Funding for open access charge: National Science Centre [grant no. 2018/31/B/NZ1/02612] and Adam Mickiewicz University.

