## Supplemental Materials for "Biochemical and genetic dissection of the RNA-binding surface of the FinO domain of *Escherichia coli* ProQ"

This pdf file includes:

Supplemental Tables S1 to S8

Supplemental Figures S1 to S15

Supplemental References

*co-corresponding authors

**SUPPLEMENTAL TABLES**

**Supplemental Table S1**. Codons identified from saturation mutagenesis screens of ProQ at positions 70 and 80 with the use of the B3H. The screen was performed with *malM* cloned into the bait construct of the assay. Both the adapter (pCW17) and bait (pKB1210) components of the assay were pre-transformed into KB473 cells in order to maximize transformation efficiency (see methods). Cells were heat shocked and plated on LB agar supplemented with inducers (0.2% arabinose and 1.5 µM IPTG), antibiotics (carbenicillin (100 µg/ml), chloramphenicol (25 µg/ml), kanamycin (50 µg/ml) and spectinomycin (100 µg/ml)) and indicators (Xgal (40 µg/ml) and TPEG (200 µM)). Plates were incubated at 37ºC overnight, then moved to 4ºC for 5-72hrs before plates were assessed for the presence of blue colonies. Blue colonies were restreaked for singles before being sent for sequencing.

| Blue colonies identified from R80X library | | |
|---|---|---|
| **Codon** | **Times isolated** | **Encodes** |
| CGT (WT) | 4 | Arg |
| AGG | 4 | Arg |
| CGC | 1 | Arg |
| CGA | 1 | Arg |
| CGG | 1 | Arg |
| AGA | 8 | Arg |
| Blue colonies identified from Y70X library | | |
| **Codon** | **Times isolated** | **Encodes** |
| TAT (WT) | 16 | Tyr |
| TAC | 3 | Tyr |

**Supplemental Table S2.** Plasmids used in the B3H study.

| **Name** | **Description** | **Details** | **Reference / Source** |
|---|---|---|---|
| pAClCI | empty vector | Encodes full-length lCI under the control of the *lauUV5* promoter; confers CamR | (Dove et al. 1997) |
| pBrα | empty vector | Encodes full-length *rpoA* under the control of tandem *lpp* and *lacUV5* promoters; confers AmpR | (Dove et al. 1997) |
| pCH1 | pCDF-1XMS2$^{hp}$ (empty vector) | pCDF-pBAD-MS2$^{hp}$-XmaI-HindIII; confers SpcR | (Pandey et al. 2020) |
| pCW17 | pAC-constitutive-λCI-MS2$^{CP}$ | Residues 1-248 of CI fused to an MS2 coat protein; transcription of this protein under the control of a constitutive promoter; confers CmR | (Pandey et al. 2020) |
| pKB949 | pBrα-ProQ$^{FL}$ | Residues 1-248 of alpha fused to full-length wild type E. coli *proQ*, serves as positive control; confers AmpR | (Pandey et al. 2020) |

| | | | |
|---|---|---|---|
| pKB955 | pBrα-ProQ$^{\Delta CTD}$ | Residues 1-248 of alpha fused to residues 2-131 of wild type E. coli proQ; confers AmpR | (Pandey et al. 2020) |
| pKB1210 | pCDF-pBAD-1xMS2$^{hp}$-malM-3'UTR | 3'UTR of *E. coli malM* (final 90 nts) cloned behind MS2hp in pCH1 between XmaI/ HindIII sites; RNA encodes its own terminator; confers SpcR | (Stein et al. 2020) |
| pSP10 | pCDF-1XMS2$^{hp}$-*cspE* 3'UTR | *E. coli* 3'UTR of *cspE* cloned behind MS2$^{hp}$ in pCH1 between XmaI/HindIII sites; mRNA encodes its own terminator; confers SpcR | (Pandey et al. 2020) |
| pSP14 | pCDF-1XMS2$^{hp}$-SibB | *E. coli sibB* cloned behind MS2$^{hp}$ in pCH1 between XmaI/HindIII sites; sRNA encodes its own terminator; confers SpcR | (Pandey et al. 2020) |
| pSP144 | pBrα-ProQ$^{\Delta CTD}$-R80K | *proQ* R80K mutation introduced to pKB955; confers AmpR | This work |
| Identified from compensatory screen: | | | |
| pSW13 | pBrα-ProQ$^{\Delta CTD}$-R80K+Y74K | *proQ* Y74K mutation introduced to pSP144; isolated from R80K-region 1 random mutagenesis library; confers AmpR | |
| pSW14 | pBrα-ProQ$^{\Delta CTD}$-R80K+Y74F | *proQ* Y74F mutation introduced to pSP144; isolated from R80K-region 1 random mutagenesis library; confers AmpR | |
| pSW15 | pBrα-ProQ$^{\Delta CTD}$-R80K+Y74L | *proQ* Y74L mutation introduced to pSP144; isolated from R80K-region 1 random mutagenesis library; confers AmpR | |
| pSW16 | pBrα-ProQ$^{\Delta CTD}$-R80K+P87N | *proQ* P87N mutation introduced to pSP144; isolated from R80K-region 2 random mutagenesis library; confers AmpR | |
| pSW17 | pBrα-ProQ$^{\Delta CTD}$-R80K+L91V | *proQ* L91V mutation introduced to pSP144; isolated from R80K-region 2 random mutagenesis library; confers AmpR | |
| pSW18 | pBrα-ProQ$^{\Delta CTD}$-R80K+L91P | *proQ* L91P mutation introduced to pSP144; isolated from R80K-region 2 random mutagenesis library; confers AmpR | |
| pSW19 | pBrα-ProQ$^{\Delta CTD}$-R80K+D92T | *proQ* D92T mutation introduced to pSP144; isolated from R80K-region 3 random mutagenesis library; confers AmpR | |
| pSW20 | pBrα-ProQ$^{\Delta CTD}$-R80K+H95R | *proQ* H95R mutation introduced to pSP144; isolated from R80K-region 3 random mutagenesis library; confers AmpR | |

**Supplemental Table S3.** Strains used in the B3H study.

| **Name** | **Relevant Details** | **Antibiotic Resistance** | **Reference/Source** |
|---|---|---|---|
| NEB 5alpha F'Iq | E. coli *lacIq* host strain for plasmid construction | TetR | New England Biolabs |
| KB473 | FW102 Δhfq cells with an F'episome which has test promoter placO$_L$2-62 fused to *lacZ* | KanR, StrR | (Berry and Hochschild 2018) |
| DH5alpha | *rec*A1, *end*A1, *lac*ZΔM15; used for plasmid amplification | AmpR | Invitrogen |

**Supplemental Table S4.** Oligonucleotides used in the B3H study.

| Name | Description | Used for | Sequence |
|---|---|---|---|
| oCG64 | F ProQ R80X library | Q5 PCR | CGGCGCAACGnnnGTCGATCTTG |
| oKB1077 | Sequencing oligo for pBrα plasmids | Sequencing | GAACAGCGTACCGACCTGG |
| oKD01 | F ProQ Y70X library | Q5 PCR | GAGCTGGCGTnnnCTTTACGGTG |
| oKD02 | R ProQ Y70X library | Q5 PCR | GAAGTGTAGAGACGTAAAGC |
| oSP92 | R ProQ R80K / R80X library | Q5 PCR | GGTTTAACACCGTAAAGATAAC |
| pSP151 | F ProQ R80K | Q5 PCR | CGGCGCAACGaagGTCGATCTTG |
| oSW8 | F ProQ-R80K region 1 library | Q5 PCR | ACCGTAAAGATAACGCCAG |
| oSW9 | R ProQ-R80K region 1 library (codon 1) | Q5 PCR | nnnAAACCCGGCGCAACGaagGTCG |
| oSW10 | R ProQ-R80K region 1 library (codon 2) | Q5 PCR | GTTnnnCCCGGCGCAACGaagGTCG |
| oSW11 | R ProQ-R80K region 1 library (codon 3) | Q5 PCR | GTTAAAnnnGGCGCAACGaagGTCG |
| oSW12 | R ProQ-R80K region 1 library (codon 4) | Q5 PCR | GTTAAACCCnnnGCAACGaagGTCG |
| oSW13 | R ProQ-R80K region 1 library (codon 5) | Q5 PCR | GTTAAACCCGGCnnnACGaagGTCG |
| oSW14 | R ProQ-R80K region 1 library (codon 6) | Q5 PCR | GTTAAACCCGGCGCAnnnaagGTCG |
| oSW15 | F ProQ region 2 library | Q5 PCR | GCCGTCAAGATCGACcttCGTTGCG |
| oSW16 | R ProQ region 2 library (codon 1) | Q5 PCR | nnnCCATGCGGTGAGCTGGACGAG C |
| oSW17 | R ProQ region 2 library (codon 2) | Q5 PCR | AACnnnTGCGGTGAGCTGGACGAG C |
| oSW18 | R ProQ region 2 library (codon 3) | Q5 PCR | AACCCAnnnGGTGAGCTGGACGAG C |
| oSW19 | R ProQ region 2 library (codon 4) | Q5 PCR | AACCCATGCnnnGAGCTGGACGAG C |
| oSW20 | R ProQ region 2 library (codon 5) | Q5 PCR | AACCCATGCGGTnnnCTGGACGAG C |
| oSW21 | R ProQ region 2 library (codon 6) | Q5 PCR | AACCCATGCGGTGAGnnnGACGAG C |
| oSW22 | F ProQ region 3 library | Q5 PCR | CAGCTCACCGCATGGGTTGC |
| oSW23 | R ProQ region 3 library (codon 1) | Q5 PCR | nnnGAGCAACATGTAGAGCATGCT CGC |
| oSW24 | R ProQ region 3 library (codon 2) | Q5 PCR | GACnnnCAACATGTAGAGCATGCT CGC |
| oSW25 | R ProQ region 3 library (codon 3) | Q5 PCR | GACGAGnnnCATGTAGAGCATGCT CGC |
| oSW26 | R ProQ region 3 library (codon 4) | Q5 PCR | GACGAGCAAnnnGTAGAGCATGCT CGC |

| | | | |
|---|---|---|---|
| oSW27 | R ProQ region 3 library (codon 5) | Q5 PCR | GACGAGCAACATnnnGAGCATGCT CGC |
| oSW28 | R ProQ region 3 library (codon 6) | Q5 PCR | GACGAGCAACATGTAnnnCATGCT CGC |
| oSW29 | R ProQ region 3 library (codon 7) | Q5 PCR | GACGAGCAACATGTAGAGnnnGCT CGC |

**Supplemental Table S5**. Details of mutagenesis libraries constructed.

| Library | Parent Plasmid | Mutagenesis oligos | Number of colonies | Theoretical # sequences | Theoretical Coverage |
|---|---|---|---|---|---|
| R80X | pCG55 | oCG64, oSP92 | 1895 | 64 | 30x |
| Y70X | pKD01 | oKD01, oKD02 | 1064 | 64 | 16x |
| R80K-region-1 | pSP144 (pKB955-R80K) | oSW8-14 | 2850 | 384 | 7x |
| R80K-region-2 | pSP144 (pKB955-R80K) | oSW15-21 | 5000 | 384 | 13x |
| R80K-region-3 | pSP144 (pKB955-R80K) | oSW22-29 | 5000 | 448 | 11x |

**Supplemental Table S6.** Plasmids used for protein overexpression.

| Name | Description | Source |
|---|---|---|
| pET-15b | empty vector; confers AmpR | Novagen |
| pET-15b-*ntd* | 1-130 aa of wild type *E. coli proQ*; confers AmpR | (Stein et al. 2020) |
| pET-15b-*ntd*_R32A | R32A mutation introduced to pET-15b-*ntd*; confers AmpR | This work |
| pET-15b-*ntd*_K35A | K35A mutation introduced to pET-15b-*ntd*; confers AmpR | This work |
| pET-15b-*ntd*_D41A | D41A mutation introduced to pET-15b-*ntd*; confers AmpR | This work |
| pET-15b-*ntd*_K54A | K54A mutation introduced to pET-15b-*ntd*; confers AmpR | This work |
| pET-15b-*ntd*_R58A | R58A mutation introduced to pET-15b-*ntd*; confers AmpR | This work |
| pET-15b-*ntd*_R62A | R62A mutation introduced to pET-15b-*ntd*; confers AmpR | This work |
| pET-15b-*ntd*_T65A | T65A mutation introduced to pET-15b-*ntd*; confers AmpR | This work |
| pET-15b-*ntd*_R69A | R69A mutation introduced to pET-15b-*ntd*; confers AmpR | This work |
| pET-15b-*ntd*_Y70F | Y70F mutation introduced to pET-15b-*ntd*; confers AmpR | This work |
| pET-15b-*ntd*_R80A | R80A mutation introduced to pET-15b-*ntd*; confers AmpR | This work |
| pET-15b-*ntd*_R80K | R80K mutation introduced to pET-15b-*ntd*; confers AmpR | This work |

**Supplemental Table S7.** Oligonucleotides used to prepare constructs for overexpression.

| **Name** | **Sequence** |
|---|---|
| R32A_F | GGAAGGTGAAGCGGCGCCGCTGAAAATCGGTATTTTT |
| R32A_R | AAAAATACCGATTTTCAGCGGCGCCGCTTCACCTTCC |
| K35A_F | GAAGCGCGTCCGCTGGCGATCGGTATTTTTCAGGATTT |
| K35A_R | AAATCCTGAAAAATACCGATCGCCAGCGGACGCGCTTC |
| D41A_F | GAAAATCGGTATTTTTCAGGCGTTGGTCGATCGTGTTG |
| D41A_R | CAACACGATCGACCAACGCCTGAAAAATACCGATTTTC |
| K54A_F | GAGCAAAACGCAATTGGCGTCCGCTTTACGTCTCTACACTT |
| K54A_R | AAGTGTAGAGACGTAAAGCGGACGCCAATTGCGTTTTGCTC |
| R58A_F | GAGCAAAACGCAATTGGCGTCCGCTTTACGTCTCTACACTT |
| R58A_R | AAGTGTAGAGACGTAAAGCGGACGCCAATTGCGTTTTGCTC |
| R62A_F | GCGATCCGCTTTAGCGCTCTACACTTCGAGCTGGCGTTAT |
| R62A_R | ATAACGCCAGCTCGAAGTGTAGAGCGCTAAAGCGGATCGC |
| T65A_F | GCTTTACGTCTCTACGCGTCGAGCTGGCGTTATC |
| T65A_R | GATAACGCCAGCTCGACGCGTAGAGACGTAAAGC |
| R69A_F | CTACACTTCGAGCTGGGCGTATCTTTACGGTGTTAAAC |
| R69A_R | GTTTAACACCGTAAAGATACGCCCAGCTCGAAGTGTAG |
| Y70F_F | CACTTCGAGCTGGCGTTTTCTTTACGGTGTTAAACCC |
| Y70F_R | GGGTTTAACACCGTAAAGAAAACGCCAGCTCGAAGTG |
| R80A_F | AAACCCGGCGCAACGGCGGTCGATCTTGACGGCAAC |
| R80A_R | GTTGCCGTCAAGATCGACCGCCGTTGCGCCGGGTTT |
| R80K_F | GTTAAACCCGGCGCAACGAAAGTCGATCTTGAC |
| R80K_R | GTCAAGATCGACTTTCGTTGCGCCGGGTTTAAC |

**Supplemental Table S8.** Oligonucleotides used to prepare templates for *in vitro* transcription.

| **Name** | **Sequence** |
|---|---|
| cspE-3′_F | TAATACGACTCACTATAGTAAGATACGTCAGCAAGAATTCAAAACC CGCTTAATC |
| cspE-3′_R | AAAAAAAACCCGCTGATTAAGCGGGTTTTGAATTCTTGCTGACGTAT CTTAC |
| cspE-3′-4U_R | AAAACCCGCTGATTAAGCGGGTTTTGAATTCTTGCTGACGTATCTTA C |
| cspE-3'-mini_F | TAATACGACTCACTATAGGAATTCGGAAGGCGCTTAATC |
| cspE-3'-mini_R | AAAAAAAAGGCGCTGATTAAGCGCCTTCCGAATTCCTAT |
| cspE-3'-mini-6U_R | AAAAAAGGCGCTGATTAAGCGCCTTCCGAATTCCTAT |
| cspE-3'-mini-4U_R | AAAAGGCGCTGATTAAGCGCCTTCCGAATTCCTAT |
| cspE-3'-mini-5'-blunt_F | TAATACGACTCACTATAGGAAGGCGCTTAATCAGCGCCTTTTTTTT |
| cspE-3'-mini-5'-blunt_R | AAAAAAAAGGCGCTGATTAAGCGCCTTCCTATAGTGAGTCG |
| cspE-3'-mini-5'-truncated_F | TAATACGACTCACTATAGGCGCTTAATCAGCGCCTTTTTTTT |
| cspE-3'-mini-5'-truncated_R | AAAAAAAAGGCGCTGATTAAGCGCCTATAGTGAGTCG |
| cspE81-3′_F | TAATACGACTCACTATAGGCCCTTCTGCTGCAAACGTAATCGCTCTG TAAGATACGTCAGCAAG |
| cspE81-3′_R | AAAAAAAACCCGCTGATTAAGCGGGTTTTGAATTCTTGCTGACGTAT CTTACAGAG |
| gapA-3′_F | TAATACGACTCACTATAGGACCTGATCGCTCACATCTCCAAATAAGT TGAGATGACACTGT |
| gapA-3′_R | AAAAAAAAGAGCGACCGAAGTCGCTCTTTTTAGATCACAGTGTCAT CTCAACTTA |
| malM-3′_F | TAATACGACTCACTATAGCTTTATCAGCAGTGTAAAAGGCAAGGGG TAATTACGCCCCACAGTG |
| malM-3′_R | AAAAAAAGGTGCGCCAGGAGACGCACCAGTTGTTGCAAAATCAGCA |

| | |
|---|---|
| | CTGTGGGGCGTAATTA |
| RybB_F | TAATACGACTCACTATAGCCACTGCTTTTCTTTGATGTCCCCATTTTG TGGAGCCC |
| RybB_R | AAAAAACCCATCAACCTTGAACCGAAATGGCGGGGTTGATGGGCTC CACAAAATGGGG |
| SibA_F | TAATACGACTCACTATAGAGGGTTAGGGAGAGGTTTCCCCCTCCCCC TGGTGTTCTTAGTAAGCCTGGAAGCTAATCACTAAGAGTA |
| SibA_R | GGGAAAGCCTCTCCCGGAGAAGAGGGCTTTTAATAAGGAAAGGGTT ATGATGAAGCACGTCATCATACTGGTGATACTCTTAGTGATTAGCT |
| SibB_F | TAATACGACTCACTATAGAGGGTAGAGCGGGGTTTCCCCCGCCCTG GTAGTCTTAGTAAGCGGGGAAGCTTATGACTAAGAGCACC |
| SibB_R | GGAAAGCCCCTCCCGAGGAAGGGGCCATAAATAAGGAAAGGGTCA TGATGAAGCTACTCATCATCGTGGTGCTCTTAGTCATAAG |
| SibB_5U_R | AAAAAGCCCCTCCCGAGGAAGGGGCCATAAATAAGGAAAGGGTCA TGATGAAGCTACTCATCATCGTGGTGCTCTTAGTCATAAG |
| SibB/cspE-3'_R | AAAAAAAAGCCTCTCCCGGAGAAGAGGGCTTTTAATAAGGAAAGGG TTATGATGAAGCACGTCATCATACTGGTGATACTCTTAGTGATTAG |

### SUPPLEMENTAL FIGURES

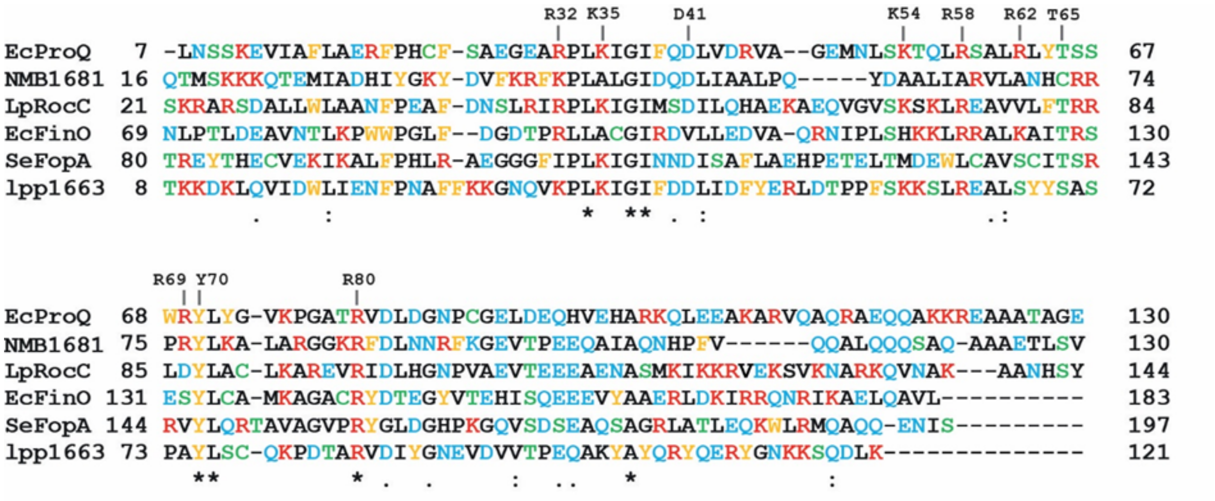


**Supplemental Figure S1. Alignment of sequences of FinO-domain proteins, which structures and/or *in vivo* RNA ligands have previously been studied.** The alignment includes the sequences of FinO domains of the following proteins: *E. coli* ProQ (Gonzalez et al. 2017; Melamed et al. 2020), *N. meningitidis* ProQ (NMB1681) (Chaulk et al. 2010; Bauriedl et al. 2020), *L. pneumophila* RocC (Attaiech et al. 2016; Kim et al. 2022), *E. coli* FinO (Ghetu et al. 2000), *S. enterica* FopA (Gerovac et al. 2020), and *L. pneumophila* lpp1663 (Attaiech et al. 2016; Immer et al. 2020). Residues whose substitutions were analyzed in this study are marked above the sequences. Positively charged residues are shown in red font, acidic residues and their amide derivatives in blue font, aromatic residues in orange font, hydroxyl or sulfhydryl group containing residues are shown in green font, and aliphatic residues in black font. Stars denote residues conserved in all six compared proteins, double dots positions where residues of the same properties are conserved, and single dots where in 5 out of 6 proteins the residues of the same properties are conserved.

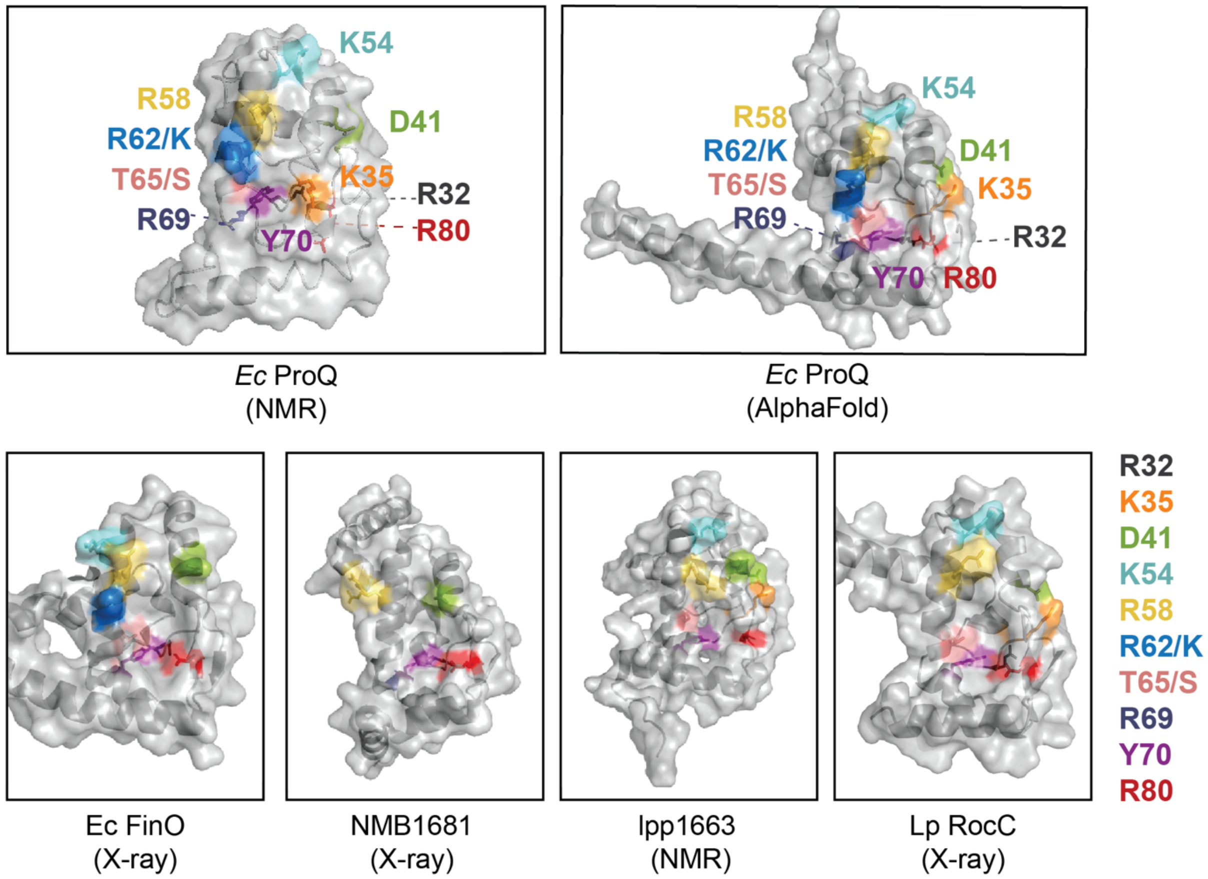


**Supplemental Figure S2. Surface representations of the structures of five FinO domains that have reported crystal or NMR structures.** The positions of the ten residues mutated in this study are each highlighted in a separate color, as indicated in the legend. Numbering is based on *E. coli* ProQ, with the homologous residues highlighted in the other proteins. FinO-domain proteins with experimentally determined structures shown here include (top left – bottom right): *E. coli* ProQ solved by NMR ((Gonzalez et al. 2017); PDB ID: 5nb9) the AlphaFold prediction for *E. coli* ProQ (Jumper et al. 2021; Varadi et al. 2022), *E. coli* F' FinO protein ((Ghetu et al. 2000); PDB ID: 1dvo), *N. meningitidis* NMB1681 ((Chaulk et al. 2010); PDB ID: 3mw6), *L. pneumophila* Lpp1663 ((Immer et al. 2020); PDB ID: 6s10), and *L. pneumophila* RocC ((Kim et al. 2022); PDB ID: 7RGU).

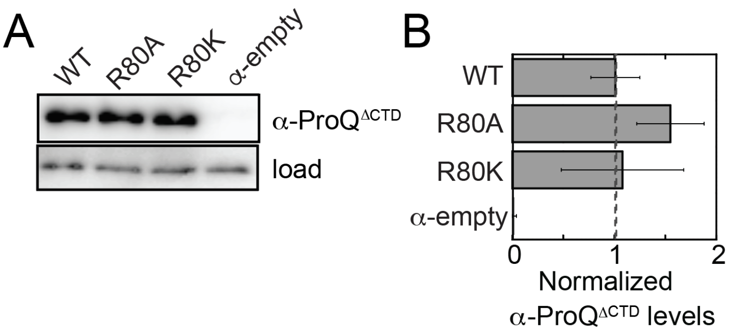


**Supplemental Figure S3. Quantification of expression of α-ProQ$^{\Delta CTD}$ variants.** (A) Samples of lysates from β-galactosidase assays from Figure 1C were analyzed by Western blot with an anti-ProQ antibody to detect the α-ProQ$^{\Delta CTD}$ fusion protein. A cross-reacting band independent of the presence of α-ProQ fusion protein is used as a loading control (load). (B) The intensity of each band (α- ProQ$^{\Delta CTD}$ and load) was quantified by densitometry and the intensity of α-ProQ$^{\Delta CTD}$ was normalized to its corresponding load band. Values for each mutant were then normalized to wild type (WT) ProQ. The bar graphs show the normalized expression levels as averages and standard deviations of values collected from at least three experiments conducted across multiple days.

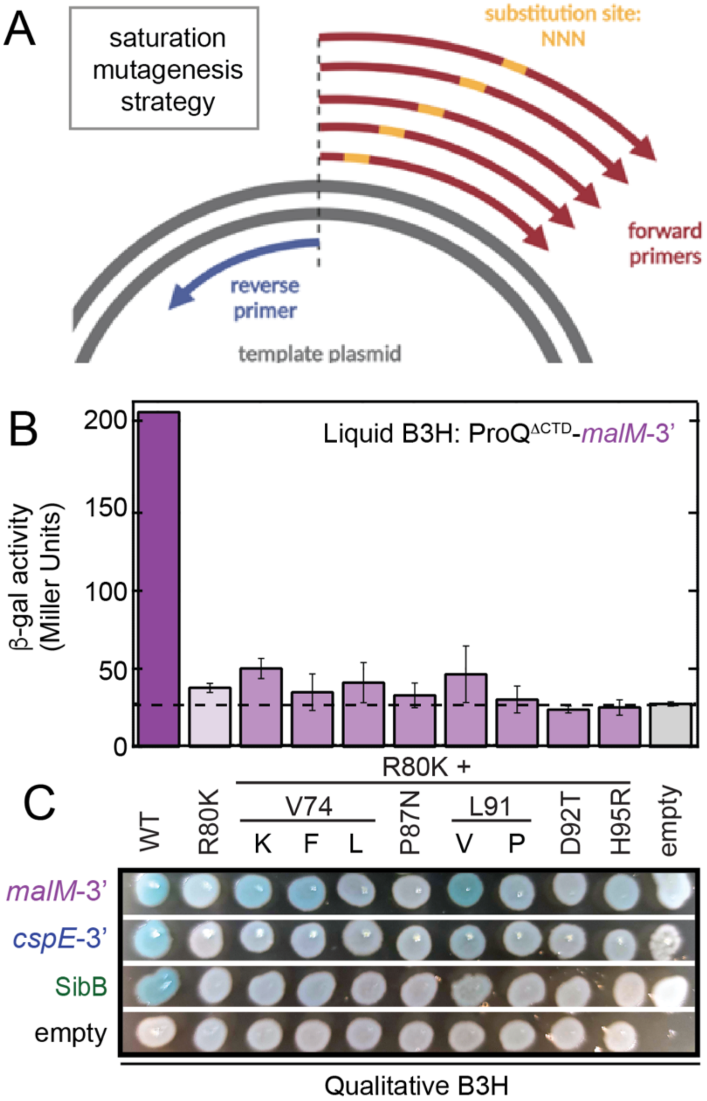


**Supplemental Figure S4. Mutagenesis strategy and results for saturation-mutagenesis screening.** (A) Schematic showing the strategy for library construction. A single reverse primer was mixed with a set of 5-7 forward primers that each contained a single randomized codon (NNN). Since each final PCR product will bear the sequence of the final primer to anneal to other intermediates, round-the-horn mutational PCR with these primers would result in PCR product containing a single amino acid substitution. (B,C) Effects of compensatory mutants across three RNA baits in B3H assay in (B) liquid assays or (C) plate-based assays. β-gal assays were performed with *Δhfq* reporter cells containing three compatible plasmids: one that encoded the CI-MS2$^{CP}$ fusion protein, another that encoded α or an α-ProQ$^{\Delta CTD}$ fusion protein (wild type, WT, or the indicated mutant), and a third that encoded a hybrid RNA (MS2$^{hp}$-*malM*-3') or an RNA that contained only the MS2$^{hp}$ moiety. The cells were grown in the presence of 0.2% arabinose and 50 μM IPTG. The bar graph shows the fold-stimulation over basal levels as averages and standard deviations of values collected from three independent experiments conducted in triplicate across multiple days.

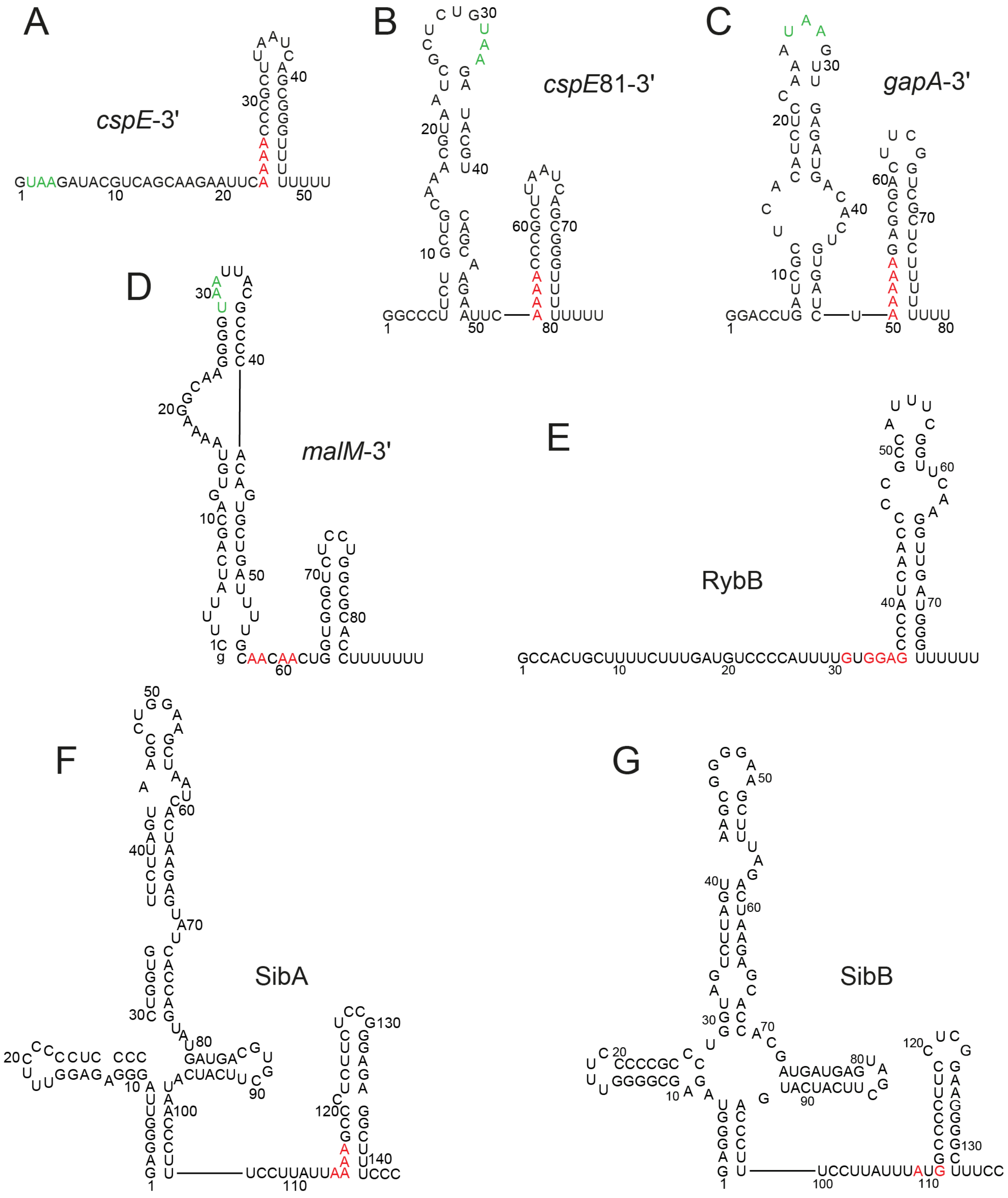


**Supplemental Figure S5. RNA ligands of ProQ protein used in this study.** The secondary structures of (A) *cspE*-3′, (B) *cspE*81, (C) *gapA*-3′, (D) *malM*-3′, and (G) SibB were predicted using *RNAstructure* software. The secondary structures of (E) RybB (Balbontín et al. 2010), and (F) SibA (Smirnov et al. 2016) are presented according to references. UAA stop codons are in green font, and the A-rich motifs located immediately 5′ of terminator hairpins are in red font. The lower case g denotes guanosine residue added on the 5' end of *malM*-3′ RNA to enable its transcription by T7 polymerase.

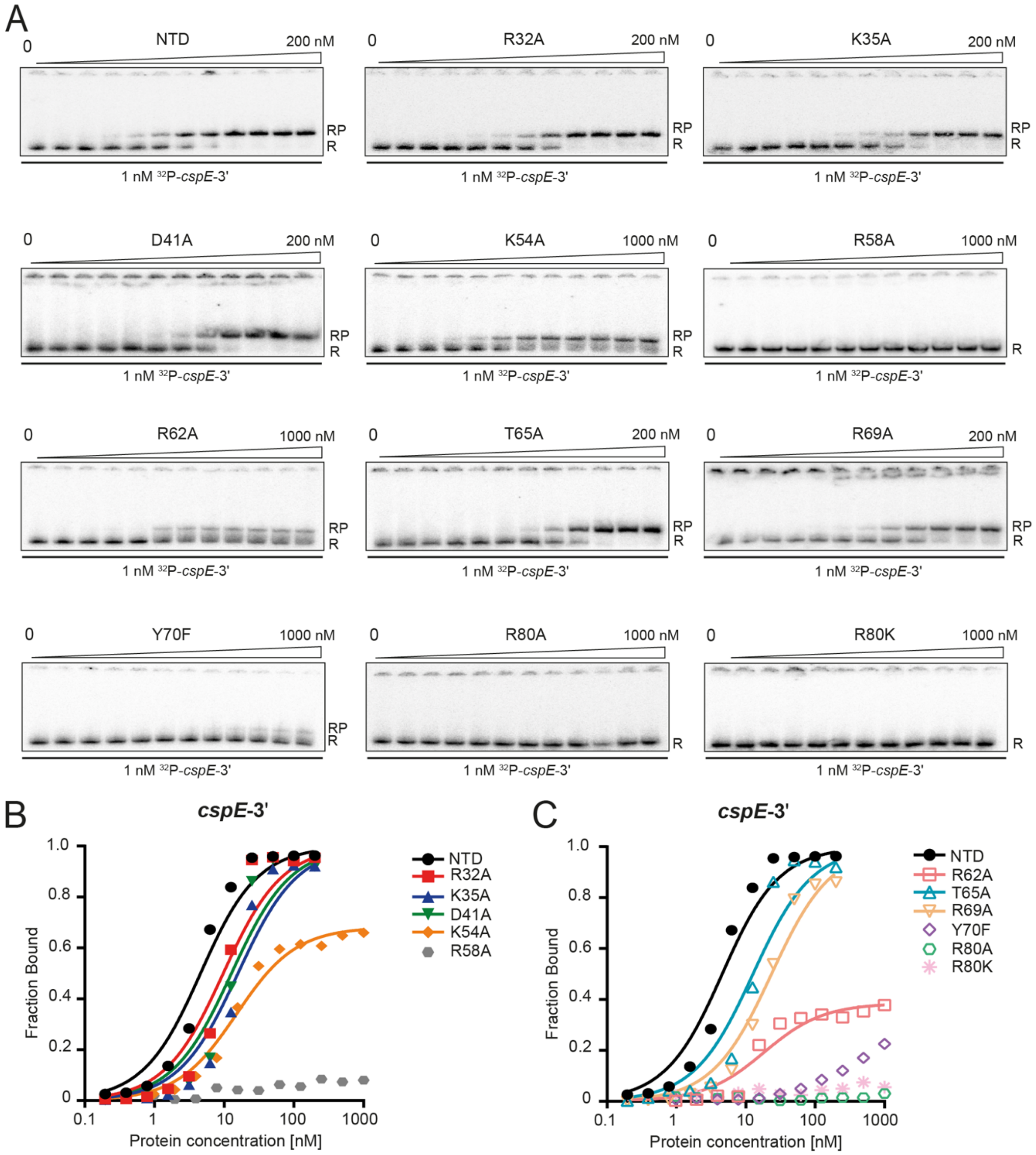


**Supplemental Figure S6. The effects of amino acid substitutions within the ProQ$^{NTD}$ (NTD) on the binding of *cspE*-3′ RNA** (A) The equilibrium binding of 1 nM $^{32}$P-*cspE*-3′ to all proteins was monitored using gelshift assay. Free $^{32}$P-RNA is marked as R, RNA-protein complexes are denoted as RP. The plots of the binding data are presented in (B) for NTD, K35A, D41A, K54A and R58A, and (C) for R62A, T65A, R69A, Y70F, R80A and R80K. The data sets in which maximum fraction bound was above 40% were analyzed by fitting to the quadratic equation, and provided $K_d$ values of 4.1 nM for NTD, 9.2 nM for R32A, 15 nM for K35A, 12 nM for D41A, 15 nM for K54A, 18 nM for R62A, 12 nM for T65A, and 23 nM for R69A. Because the maximum fraction bound of *cspE*-3' to Y70F mutant was below 40%, the $K_d$ value was assumed as bigger than 1μM. The binding of *cspE*-3' to R58A, R80A and R80K mutants was essentially undetectable up to 1μM concentrations. Average equilibrium dissociation constant ($K_d$) values for RNA binding to ProQ are shown in the Table 1 (main text).

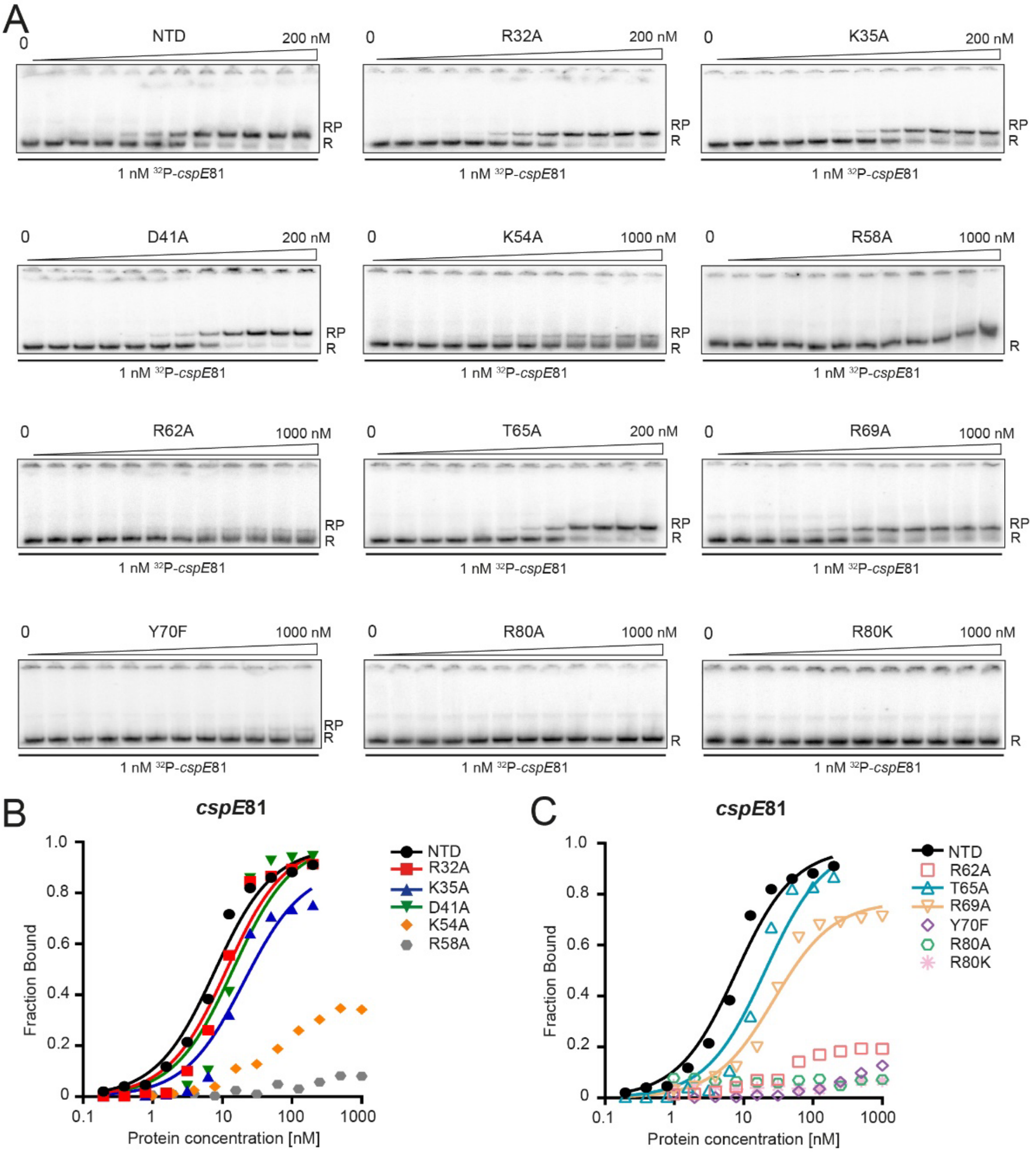


**Supplemental Figure S7. The effects of amino acid substitutions within the ProQ$^{NTD}$ (NTD) on the binding of *cspE*81 RNA.** (A) The equilibrium binding of 1 nM $^{32}$P-*cspE*81 to all proteins was monitored using gelshift assay. Free $^{32}$P-RNA is marked as R, RNA-protein complexes are denoted as RP. The plots of the binding data are presented in (B) for NTD, K35A, D41A, K54A and R58A, and (C) for R62A, T65A, R69A, Y70F, R80A and R80K. The data sets in which maximum fraction bound was above 40% were analyzed by fitting to the quadratic equation, and provided $K_d$ values of 7.2 nM for NTD, 11 nM for R32A, 20 nM for K35A, 14 nM for D41A, 20 nM for T65A, and 27 nM for R69A. Because the maximum fractions bound of *cspE*81 to K54A, R62A and Y70F mutants were below 40%, the $K_d$ values were assumed as bigger than 1μM. The binding of *cspE*81 to R58A, R80A and R80K mutants was essentially undetectable up to 1μM concentrations. Average equilibrium dissociation constant ($K_d$) values for RNA binding to ProQ are shown in the Table 1 (main text).

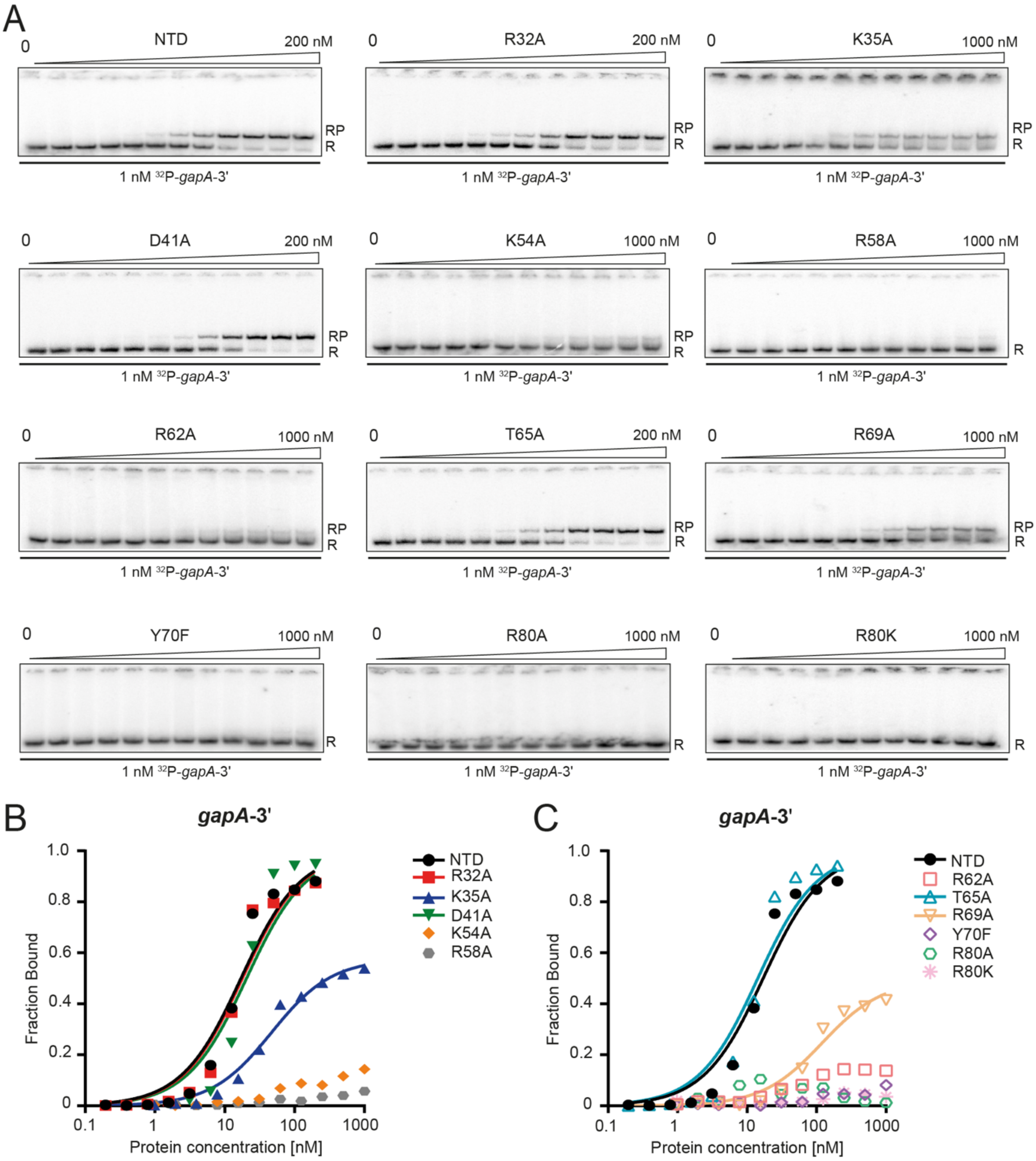


**Supplemental Figure S8. The effects of amino acid substitutions within the ProQ$^{NTD}$ (NTD) on the binding of *gapA*-3' RNA.** (A) The equilibrium binding of 1 nM $^{32}$P- *gapA*-3' to all proteins was monitored using gelshift assay. Free $^{32}$P-RNA is marked as R, RNA-protein complexes are denoted as RP. The plots of the binding data are presented in (B) for NTD, K35A, D41A, K54A and R58A, and (C) for R62A, T65A, R69A, Y70F, R80A and R80K. The data sets in which maximum fraction bound was above 40% were analyzed by fitting to the quadratic equation, and provided $K_d$ values of 17 nM for NTD, 17 nM for R32A, 46 nM for K35A, 20 nM for D41A, 14 nM for T65A, and 120 nM for R69A. Because the maximum fractions bound of *gapA*-3' to K54A and R62A mutants were below 40%, the $K_d$ values were assumed as bigger than 1µM. The binding of *gapA*-3' to R58A, Y70F, R80A and R80K mutants was essentially undetectable up to 1µM concentrations. Average equilibrium dissociation constant ($K_d$) values for RNA binding to ProQ are shown in the Table 1 (main text).

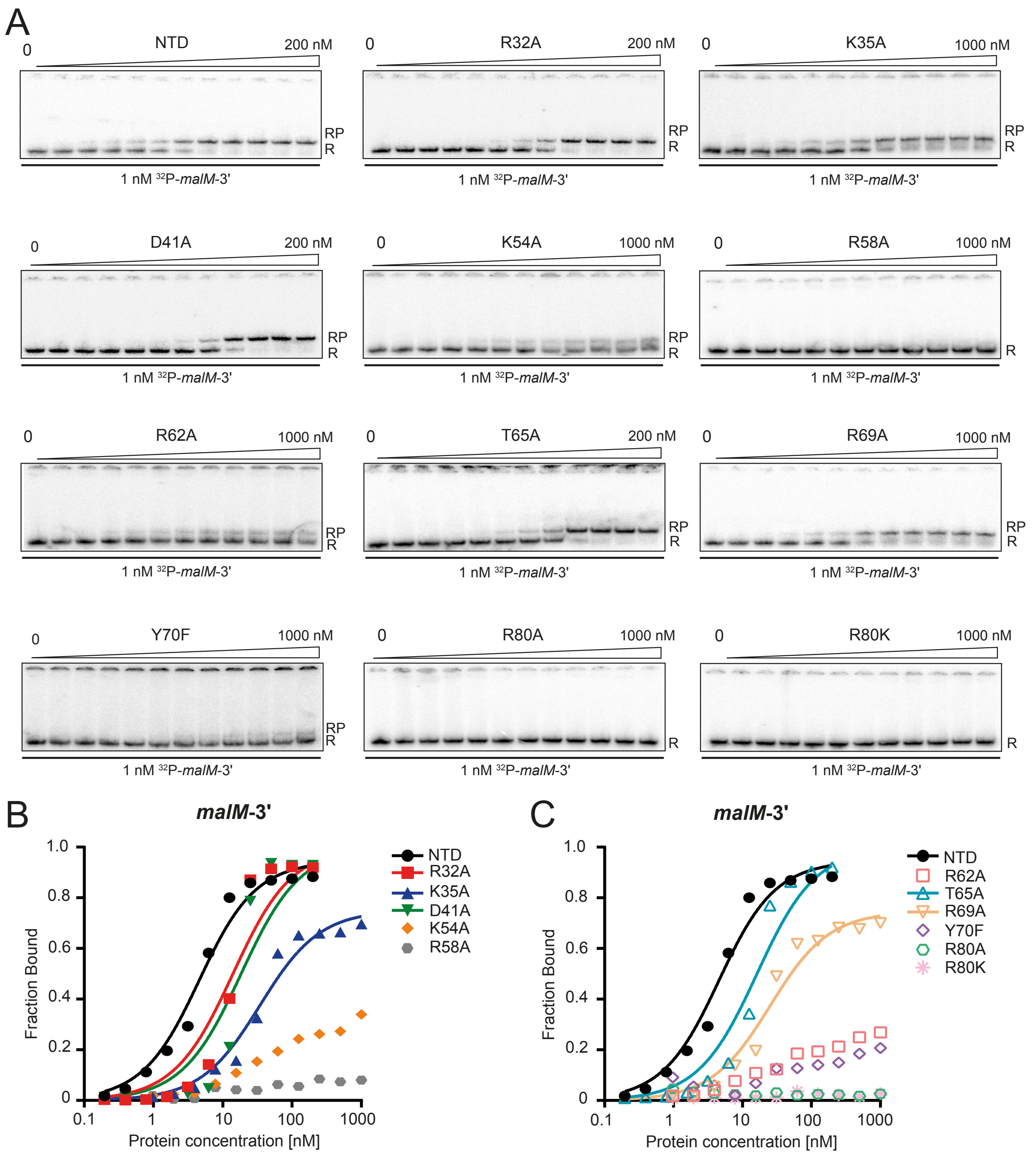


**Supplement Figure S9. The effects of amino acid substitutions within the ProQ$^{NTD}$ (NTD) on the binding of *malM*-3′ RNA**. (A) The equilibrium binding of 1 nM $^{32}$P-*malM*-3′ to all proteins was monitored using gelshift assay. Free $^{32}$P-RNA is marked as R, RNA-protein complexes are denoted as RP. The plots of the binding data are presented in (B) for NTD, K35A, D41A, K54A and R58A, and (C) for R62A, T65A, R69A, Y70F, R80A and R80K. The data sets in which maximum fraction bound was above 40% were analyzed by fitting to the quadratic equation, and provided $K_d$ values of 4.1 nM for NTD, 14 nM for R32A, 34 nM for K35A, 18 for D41A, 16 nM for T65A, and 25 nM for R69A. Because the maximum fractions bound of K54A, R62A and Y70F mutants were below 40%, the $K_d$ values were assumed as bigger than 1µM. The binding of *malM*-3' to R58A, R80A and R80K mutants was essentially undetectable up to 1µM concentrations. Average equilibrium dissociation constant ($K_d$) values for RNA binding to the NTD are shown in the Table 1 (main text).

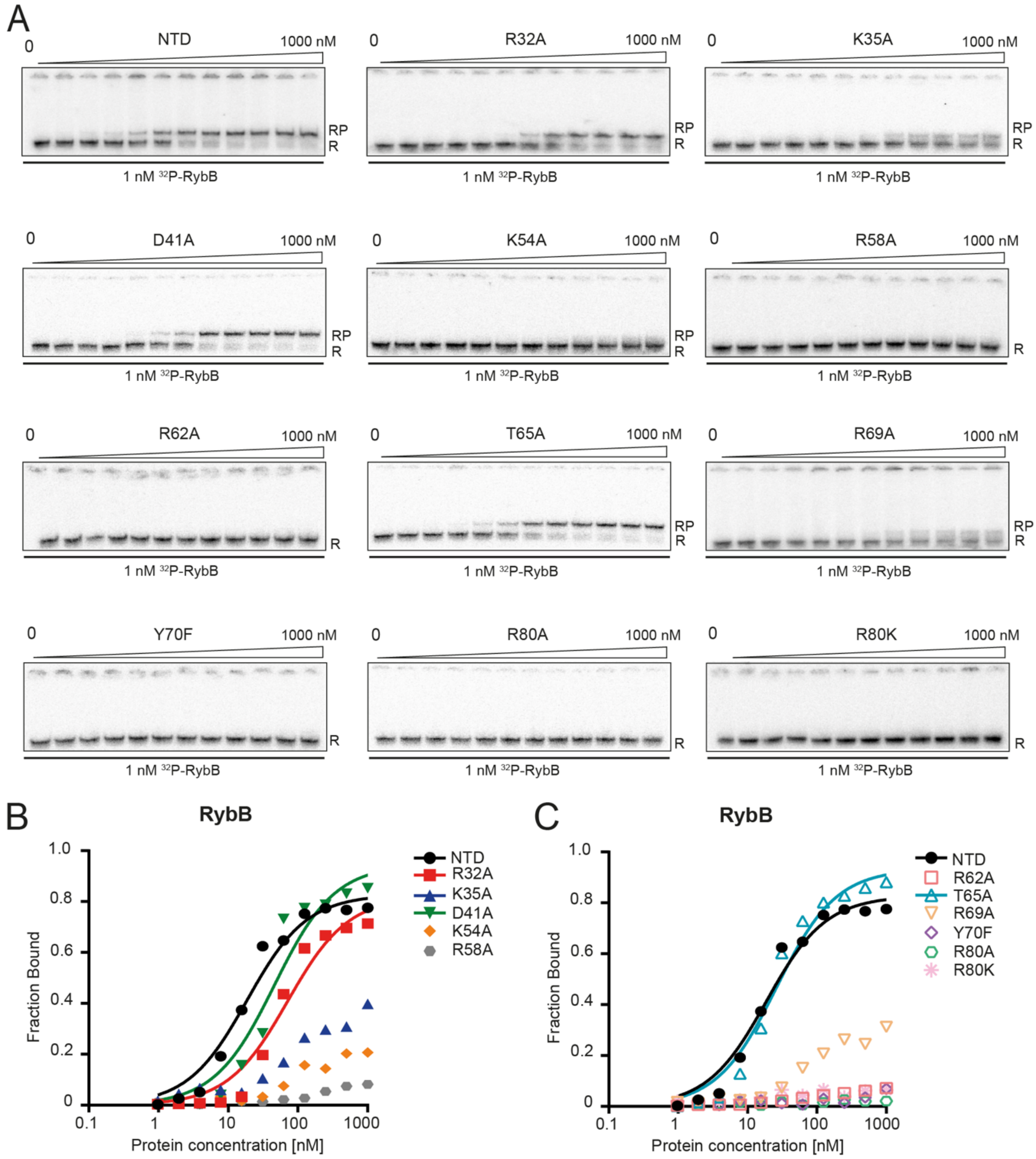


**Supplemental Figure S10. The effects of amino acid substitutions within the ProQ$^{NTD}$ (NTD) on the binding of RybB RNA.** (A) The equilibrium binding of 1 nM $^{32}$P-RybB to all proteins was monitored using gelshift assay. Free $^{32}$P-RNA is marked as R, RNA-protein complexes are denoted as RP. The plots of the binding data are presented in (B) for NTD, K35A, D41A, K54A and R58A, and (C) for R62A, T65A, R69A, Y70F, R80A and R80K. The data sets in which maximum fraction bound was above 40% were analyzed by fitting to the quadratic equation, and provided $K_d$ values of 18 nM for NTD, 71 nM for R32A, 46 nM for D41A, and 26 nM for T65A. Because the maximum fractions bound of RybB to K35A, K54A, and R69A mutants were below 40%, the $K_d$ values were assumed as bigger than 1μM. The binding of RybB to R58A, R62A, Y70F, R80A and R80K mutants was essentially undetectable up to 1μM concentrations. Average equilibrium dissociation constant ($K_d$) values for RNA binding to ProQ are shown in the Table 1 (main text).

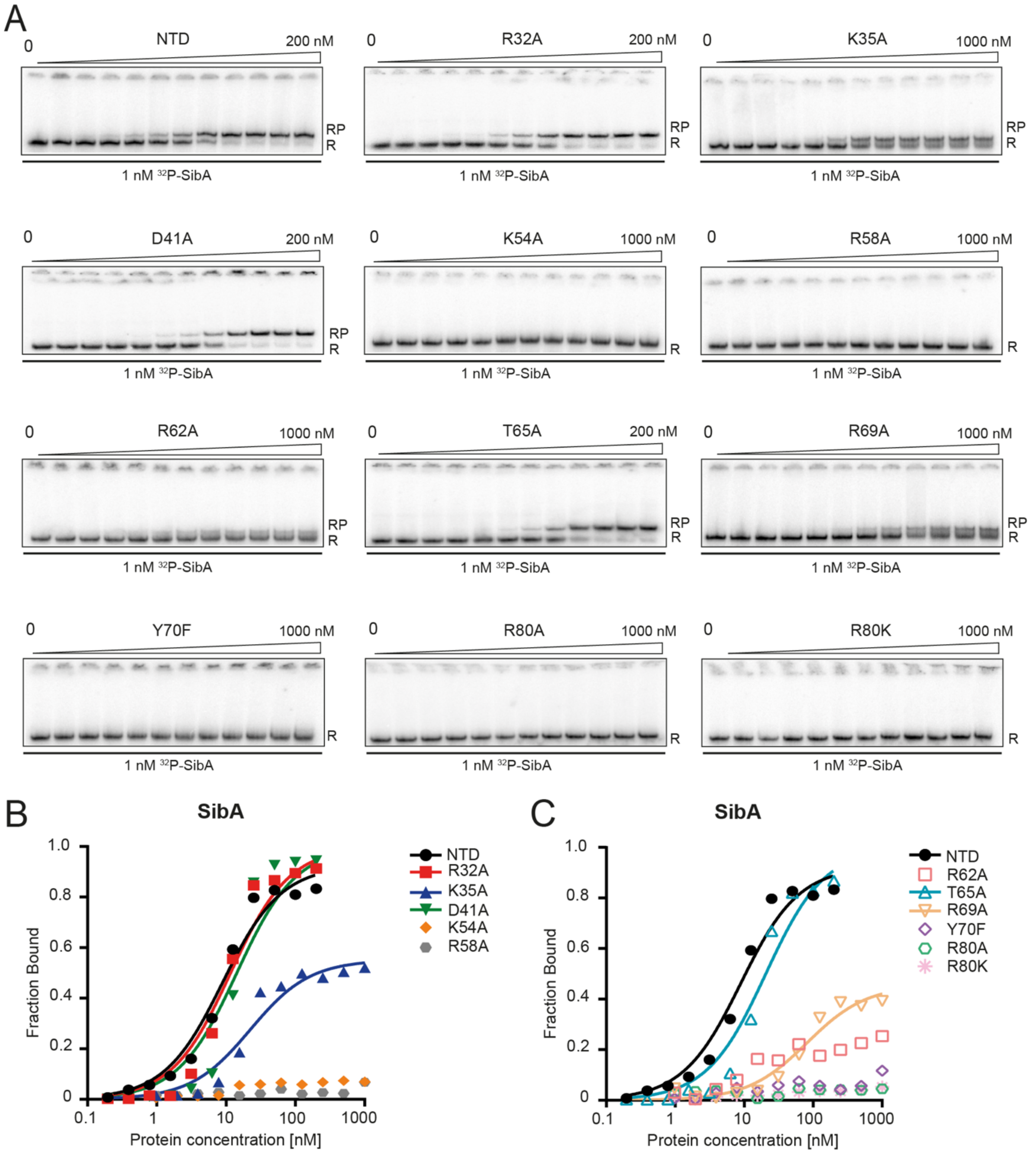


**Supplemental Figure S11. The effects of amino acid substitutions within the ProQ$^{NTD}$ (NTD) on the binding of SibA RNA.** (A) The equilibrium binding of 1 nM $^{32}$P-SibA to all proteins was monitored using gelshift assay. Free $^{32}$P-RNA is marked as R, RNA-protein complexes are denoted as RP. The plots of the binding data are presented in (B) for NTD, K35A, D41A, K54A and R58A, and (C) for R62A, T65A, R69A, Y70F, R80A and R80K. The data sets in which maximum fraction bound was above 40% were analyzed by fitting to the quadratic equation, and provided $K_d$ values of 8.4 nM for NTD, 11 nM for R32A, 22 nM for K35A, 14 nM for D41A, 20 nM for T65A, and 81 nM for R69A. Because the maximum fraction bound of SibA to R62A mutant was below 40%, the $K_d$ value was assumed to be bigger than 1μM. The binding of SibA to K54A, R58A, Y70F, R80A and R80K mutants was essentially undetectable up to 1μM concentrations. Average equilibrium dissociation constant ($K_d$) values for RNA binding to ProQ are shown in the Table 1 (main text).

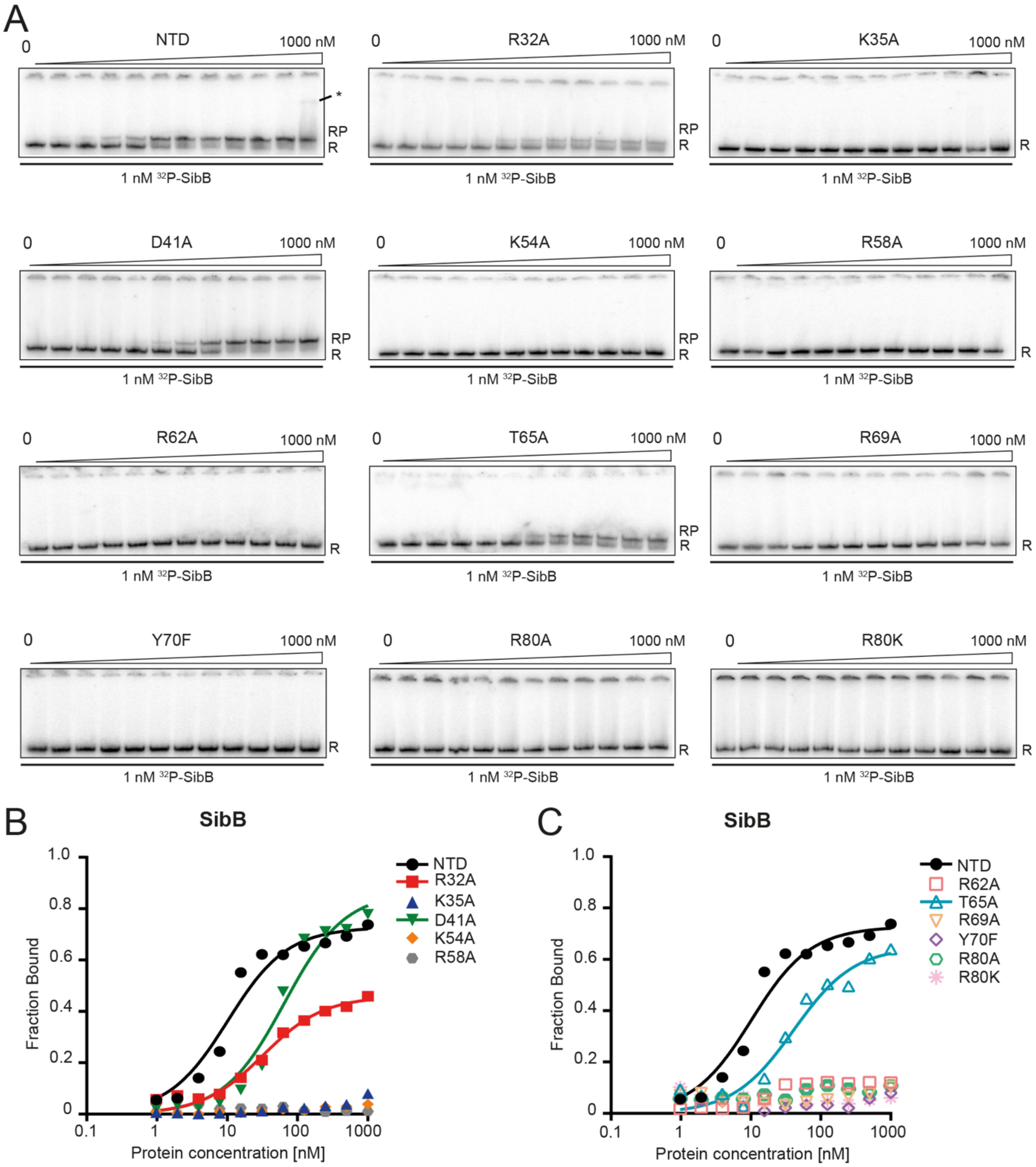


**Supplemental Figure S12. The effects of amino acid substitutions within the ProQ$^{NTD}$ (NTD) on the binding of SibB RNA.** (A) The equilibrium binding of 1 nM $^{32}$P-SibB to all proteins was monitored using gelshift assay. Free $^{32}$P-RNA is marked as R, RNA-protein complexes are denoted as RP. The plots of the binding data are presented in B) for NTD, K35A, D41A, K54A and R58A, and (C) for R62A, T65A, R69A, Y70F, R80A and R80K. The data sets in which maximum fraction bound was above 40% were analyzed by fitting to the quadratic equation, and provided $K_d$ values of 9.7 nM for NTD, 32 nM for R32A, 64 nM for D41A and 40 nM for T65A. The binding of SibA to K35A, K54A, R58A, R62A, R69A, Y70F, R80A and R80K mutants was essentially undetectable up to 1μM concentrations. Average equilibrium dissociation constant ($K_d$) values for RNA binding to ProQ are shown in Table 1 (main text).

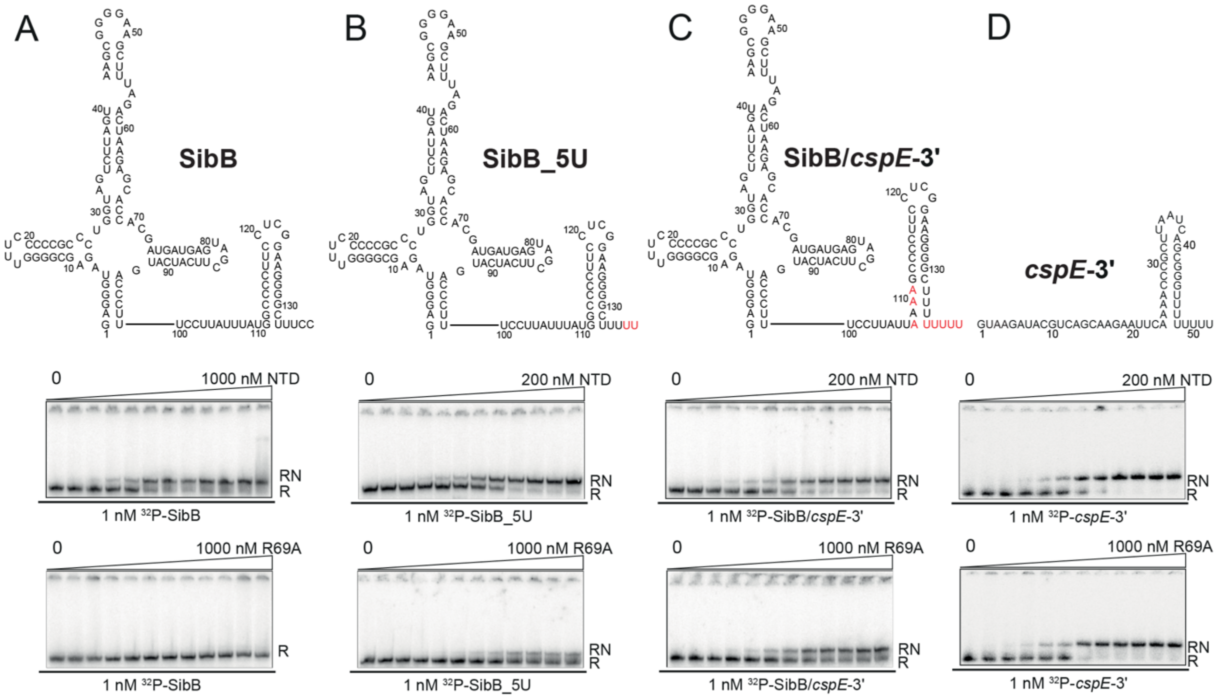


**Supplemental Figure S13. Raw gel data, which show the effects of mutations in the terminator hairpin of SibB on its binding to WT ProQ$^{NTD}$ and its R69A mutant.** The plots of the presented data are shown in Figure 5, and the calculated $K_d$ values in the Table 2 of the main text.

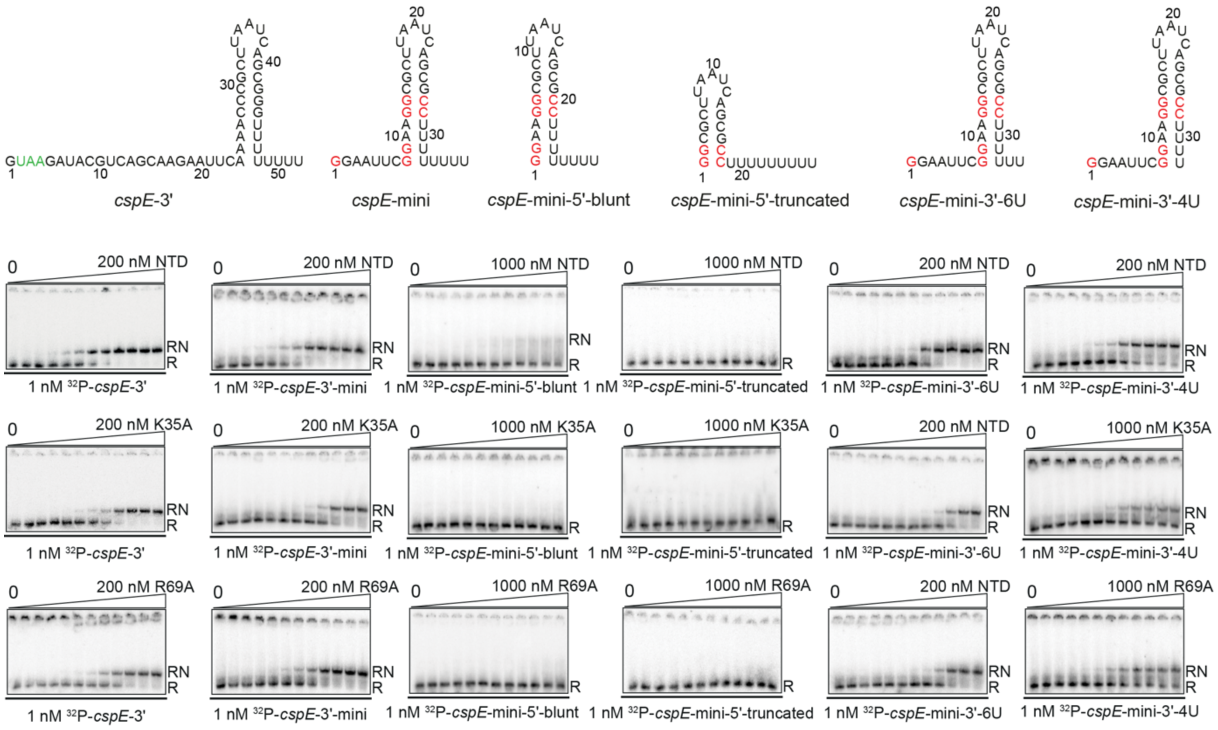


**Supplemental Figure S14. Gelshift analysis of the binding of *cspE*-3′, *cspE*-mini, *cspE*-mini-5′-blunt, *cspE*-mini-5′-truncated, *cspE*-mini-3′-6U, and *cspE*-mini-3′-4U to the WT ProQ$^{NTD}$ (NTD), and its K35A and R69A mutants.** Free 1 nM $^{32}$P-RNA is marked as R, RNA-protein complexes are denoted as RP. The fitting of the data represented on gels, and the $K_d$ values are shown in Fig. 6, and Table 2 in the main text.

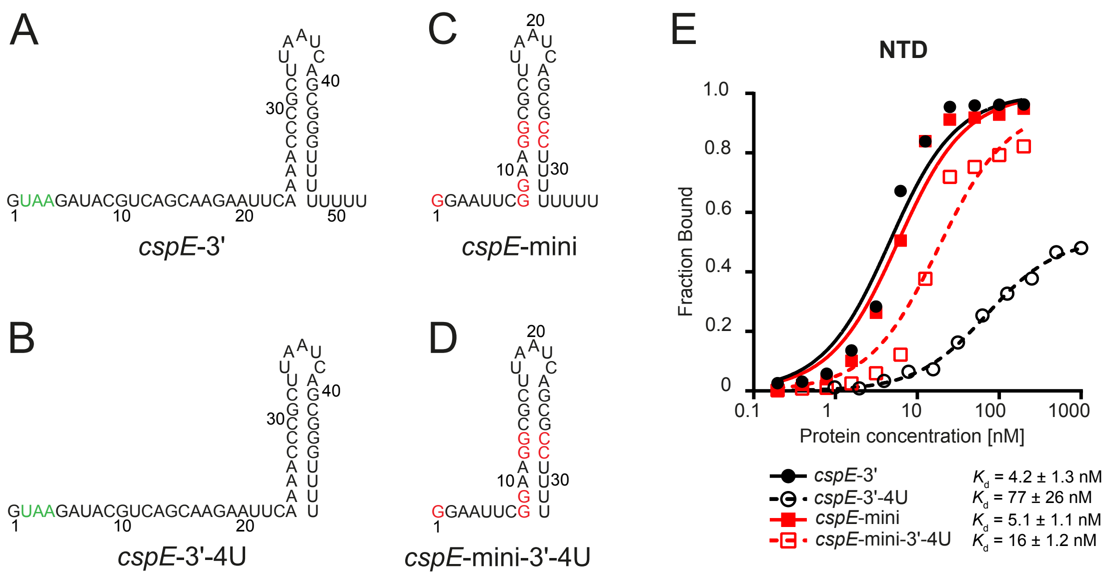


**Supplemental Figure S15. Different effects of the truncation of the 3'-oligoU-tail in *cspE*-3'-4U and *cspE*-mini-3′-4U molecules.** Equilibrium binding of 1 nM (A) $^{32}$P-*cspE*-3′, (B) $^{32}$P-*cspE*-3′-4U, (C) $^{32}$P-*cspE*-mini, and (D) $^{32}$P-*cspE*-mini-4U to the NTD was monitored using gelshift assay. The NTD concentration series was prepared by 2-fold sequential dilutions in the concentration range 0-200 nM for *cspE*-3′, *cspE*-mini and *cspE*-mini-4U, and 0-1000 nM for *cspE*-3′-4U. (E) The fitting of the binding data was performed using the quadratic equation. Data for *cspE*-3′, *cspE*-mini and *cspE*-mini-4U binding to WT ProQ$^{NTD}$ are the same as in Figure 6 (main text) and Suppl. Fig. S14. Average equilibrium dissociation constant ($K_d$) values with standard deviation from at least three independent experiments are shown in the legend.
